# The Glycoproteome of *Loxosceles* Venom Reveals an Expanded Landscape of Animal Toxins

**DOI:** 10.64898/2026.10.29.762871

**Authors:** Raniele S. Moreira, Bruno C. S. Silva, Dener E. Bortolini, Manuela C. E. Ferreira, Carlena T. N. de Reyes, Marcele N. Rocha, Milene B. Carvalho, Martin R. Larsen, Carlos Chávez-Olórtegui, Franck Molina, Laurence Molina, João C. Minozzo, Marcella N. Melo-Braga, Liza F. Felicori

## Abstract

Spider venoms of the genus *Loxosceles* constitute a complex mixture of bioactive proteins and peptides associated with the major clinical manifestations of loxoscelism. Although significant advances have been made in the proteomic characterization of these toxins, studies addressing post-translational modifications, particularly glycosylation, and their potential impact on venom properties remain scarce. Here, we investigated the occurrence and characteristics of N-glycosylation in the venoms of *Loxosceles gaucho*, *Loxosceles intermedia*, and *Loxosceles laeta*. Using two-dimensional electrophoresis combined with glycoprotein-specific staining and LC-MS/MS, we identified glycosylation in different proteins biologically relevant to envenomation, predominantly belonging to the phospholipase D family, with distinct profiles among the species analyzed. Bottom-up glycoproteomic analysis of crude venoms using Byonic and a customized sequence database identified 88 N-glycosylated peptide sequences, primarily associated with lyases and hydrolases.

Extensive microheterogeneity was detected at individual glycosylation sites, with paucimannosidic structures predominating alongside oligomannose and hybrid/complex N-glycans. Complementary enzyme-linked lectin assays supported the presence of distinct carbohydrate epitopes, including mannose, galactose, and N-acetylgalactosamine containing structures, and revealed α2,6-sialic acid-reactive motifs in two species. These findings establish N-glycosylation as an additional layer of molecular heterogeneity in *Loxosceles* venoms and provide a foundation for investigating its potential contribution to toxin properties and venom diversity.

## Introduction

Spiders of the genus *Loxosceles* have a global distribution and are responsible for accidents that can cause severe clinical manifestations, known as loxoscelism. In South America, the species *L. intermedia*, *L. gaucho*, and *L. laeta* are of greatest medical importance ^1^. The pathophysiology of envenomation is related to the various components of the venom, which is composed predominantly of proteins. Biochemical analyses using one-or two-dimensional electrophoresis associated with mass spectrometry have shown that crude venoms contain proteins primarily concentrated in the 2-40 kDa range. However, molecules with molecular masses above this interval are also detected, though in lower proportions ^2–5^.

Transcriptomic studies of the venom glands of these species reveal an abundance of Phospholipases D (PLDs) (30-35 kDa) and metalloproteases (20-30 kDa), with the former being the main toxin family involved in the effects of envenomation and consequently the most thoroughly characterized ^6,7^. In lower proportions, but also associated with documented effects in human envenomation, hyaluronidases (40-50 kDa), serine proteases (90-95 kDa), among others, have been identified ^8–11^.

The numerous studies aimed at elucidating the structural and biological aspects of *Loxosceles* venom make evident both the importance of its toxins and their complex nature, acting as they do on diverse targets. Nevertheless, certain molecular aspects of the venom remain poorly explored, such as post-translational modifications (PTMs).

Among the more than 300 PTMs known to occur physiologically, glycosylation stands out for its frequency and importance in structural and functional processes of proteins ^12–15^. Despite this, data on this modification in arthropod venoms remain limited. In a previous study by our group, it was found that 83% of glycosylation records in arthropod toxins were putative. Among these toxins, the PLDs, which are abundant in the *Loxosceles* venoms, had the highest number of putative N-glycosylation sites ^16^.

N-glycosylation, the focus of this work, involves the covalent attachment of glycans to the amide nitrogen of asparagine residues within the consensus sequence N-X-S/T, where X is any amino acid except proline ^17^. However, importantly, the presence of this consensus sequence does not necessarily indicate that a site is glycosylated, as site occupancy depends on protein structure and the cellular glycosylation machinery.

There is already functional evidence suggesting that glycosylation is relevant in dermonecrotic activity of spider *Loxosceles intermedia* venom ^18^.

However, the glycosylated proteins were neither identified nor characterized, leaving their potential biological roles unclear.

Advances in glycoproteomics now enable the identification of glycosylated proteins, localization of glycosylation sites, and characterization of the associated glycan structures. These approaches therefore provide an opportunity to move beyond the identification of putative glycosylation sites and determine which sites are actually occupied in venom proteins. Given the potential relevance of glycosylation to venom protein function and the limited experimental evidence available for *Loxosceles* venoms, this study aimed to identify and characterize glycosylation in the venoms of the medically important species *L. intermedia*, *L. gaucho*, and *L. laeta*.

## Materials and methods

### 1.1 Venom

Whole venom from *Loxosceles intermedia*, *L. laeta*, and *L. gaucho* were provided by CPPI (Center for the Production and Research on Immunobiologicals; Curitiba, Paraná, Brazil). Venoms were collected by applying a 15 V electrical stimulus to the cephalothorax of the spiders. Each venom batch consisted of pooled venom collected from a large population of both male and female spiders, comprising thousands of individuals. The venom batches used throughout this study were SVLICPPI 128/21 (*L. intermedia*), SVLGCPPI 120/18 (*L. gaucho*), and SVLLCPPI 1221/18 (*L. laeta*).

### 1.2 Two-dimensional gel electrophoresis (2DE)

Protein venom concentration was determined using a Qubit Protein Assay Kit, and 60 μg of protein from each venom was precipitated in acetone at a 1:6 ratio. All experiments were performed in three independent technical replicates. After overnight precipitation, samples were centrifuged at 15,000g, 4°C for 10 minutes. The supernatant was removed, and the pellet resuspended in 125 μL of 2DE rehydration buffer (7 M urea, 2 M thiourea, 2% CHAPS, 0.5% IPG buffer, pH 3–10, 1% protease inhibitor, 0.002% Bromophenol blue and 18 mM DTT). Venom samples were applied to a 7 cm immobilized pH gradient (IPG) strip in a non-linear range of pH 3–10 (GE Healthcare) and subjected to passive rehydration overnight at room temperature. Isoelectric focusing (IEF) was performed at 20° C on an IPGphor unit (IPGphor 3 Isoelectric Focusing System – GE Healthcare) with a total voltage accumulation of 15,375 V for 9h (Step 500 V/3h, Step 1000/30min, Gradient 5000/2h30min, Step 5000V/1h, Step 500V/2h) at 50 μA/strip. After focusing, the strips were incubated for 15 min in reducing solution (6 M urea, 30% glycerol, 2% SDS, 75 mM Tris-HCl pH 8.8 and 64.82 mM DTT), followed by a 15 min incubation in alkylation solution (6 M urea, 30% glycerol, 2% SDS, 75 mM Tris-HCl pH 8.8 and 135.16 mM iodoacetamide). The sodium dodecyl sulfate polyacrylamide gel electrophoresis (SDS-PAGE) step was performed using 12.5% polyacrylamide gels at constant voltage of 100V. Glycosylated proteins were detected using the Pro-Q® Emerald 300 Glycoprotein Gel and Blot Stain kit (Thermo Scientifics) and total proteins were detected using SYPRO ® Ruby protein gel stain, according to the manufacturer’s instructions. Gel spot detection and analysis were performed using Image Master 2D Platinum 6.0 software (GE Healthcare).

### 1.3 In gel Digestion

For mass spectrometry analysis, venom protein samples were separated by 2DE and stained with Coomassie blue. Spots consistently detected across independent 2DE replicates were selected for LC-MS/MS. Representative spots displaying strong glycoprotein staining and multiple proteoforms with similar molecular masses but distinct isoelectric points were prioritized, with preference given to those within the molecular weight range showing the highest frequency of glycosylation. The process was performed as previously described by Schevchenko et al ^19^ with minor modifications. Eleven spots (4 from *L. intermedia*, 4 from *L. gaucho*, and 3 from *L. laeta* gels) were manually excised and destained with washes of ethanol and Mili-Q water. The proteins were reduced with 10 mM dithiothreitol at 56 °C for 45 minutes and alkylated with 55 mM iodoacetamide (IAA) and digested with proteomics-grade trypsin (PROMEGA) in 50 mM ammonium bicarbonate (TEAB). After 45 minutes, the supernatant was removed and 50 mM ammonium bicarbonate (TEAB) pH 8.3 was added to the gel overnight at 37 °C and stopped by adding 5% formic acid. The peptides in gel were extracted using a solution containing 5% formic acid and 20% acetonitrile. The peptides were vacuum-dried, resuspended in 0.1% formic acid, and cleaned-up using Ziptip C18 according to manufacture instructions. The eluted samples were dried in the Speed Vac.

### 1.4 The mass spectrometry analysis of the spots

After in-gel digestion, peptides were resuspended in 0.1 % formic acid and analyzed by LC-MS/MS using a Thermo Easy-nLC system coupled to an Orbitrap Lumos mass spectrometer with a 55 min gradient. MS spectra were acquired in the Orbitrap (resolution 120,000; m/z 350–1500; maximum injection time 50 ms). MS/MS acquisition was performed in data-dependent mode (cycle time) at a resolution of 50,000 using HCD fragmentation (CE 32) and a maximum injection time of 200 ms. Raw data were processed in Byonic (Protein Metrics Suite v5.12) for protein and glycopeptide identification against a Custom *Loxosceles* Protein Database described below. Searches were performed with trypsin (up to two missed cleavages), carbamidomethylation of cysteine as a fixed modification, and methionine oxidation and N-terminal pyroglutamate formation as variable modifications. N-linked glycosylation on asparagine residues was evaluated using a database of 38 insect-type N-glycans. Precursor and fragment mass tolerances were set to 10 ppm and 0.05 Da, respectively. Identifications were filtered to a false discovery rate (FDR) < 1%. Only glycopeptides with 2D posterior error probability (PEP-2D) < 0.001 and Byonic scores > 100 were retained ^20^. The glycan compositions assigned by Byonic were used to infer the corresponding N-glycan structures using GlycoWorkbench (version 1.0.3353). Glycan structures were manually assembled according to the identified monosaccharide compositions and structural features commonly reported for insect glycoproteins and venoms. Due to the limitations of the analytical approach, the exact position of specific monosaccharide residues, glycan branching, and α-or β-glycosidic linkages could not be experimentally determined.

### 1.5 Generation of a Custom *Loxosceles* spp. Protein Database

The transcriptomes of *L. laeta* (EY188373–EY189729) and *L. intermedia* (HO002618–HO004460) from GenBank, as well as the coding DNA sequences (CDS) of the L. reclusa genome (Brown Recluse Spider Genome Project, Baylor College of Medicine) were downloaded. As the *L. intermedia* transcriptome was composed by Expressed sequence tags (ESTs), the sequences were assembled using CAP3 software ^21^ with an overlap length cutoff of 40 bp., a minimum overlap identity of 97%, and default parameters for all remaining options, resulting in 575 sequences (207 contigs and 368 singlets). Genomic and transcriptomic sequences from species of the genus *Loxosceles* were retrieved from GenBank, GenPept, UniProt, and Arachno Server public databases with searches conducted between 2019 and 2020 (DOI: 10.5281/zenodo.22876588).

DNA and RNA sequences were translated into their corresponding amino acid sequences using an in-house-developed Python 3 script ^22^ that utilized the Biopython library ^23^. This script selects, for each sequence, the longest open reading frame (ORF) bounded either by stop codons or by ambiguous residues (“X”) resulting from the translation of undefined nucleotides (“N”). Redundant sequences were then filtered using CD-HIT Suite ^24^ with a sequence identity cutoff of 99.9%. Sequences obtained in the previous steps were compiled into a FASTA file for use in the glycoproteomic analysis and comprise the Protein *Loxosceles* spp. Sequence Database.

To perform functional annotation of the sequences in the Custom *Loxosceles* spp. Protein Database based on their similarity to publicly available sequences from NCBI (GenBank, RefSeq, TPA, SwissProt, PIR, PRF, and PDB), UniProt, and ArachnoServer, nucleotide and amino acid databases were independently generated for the family Sicariidae and the suborder Araneomorphae, resulting in a total of four databases. Unannotated sequences were first aligned against the Sicariidae databases, and sequences that remained unannotated were subsequently aligned against the Araneomorphae databases. BLASTp, BLASTn, and tBLASTx were used for these alignments according to the type of target database. The remaining unannotated sequences were aligned against the nr database using DIAMOND BLASTx. Only alignments with an e-value < 1.0 × 10⁻⁵ (1.0 × 10⁻¹⁰ for tBLASTx), query coverage > 70%, and bit score > 50 were considered for annotation, and only the top-ranking hit (best hit) was considered for each sequence. For the L. reclusa genome, the second alignment step was performed against the Nucleotide Collection (nt) database instead of the Araneomorphae database. Despite the alignments performed against these databases, some sequences in the Custom *Loxosceles* spp. Protein Database remained unannotated.

### 1.6 Glycoproteomic analysis

Bottom-up glycoproteomics was performed as previously described by Verano-Braga et al. ^25^, with minor modifications. Approximately 500 µg of venom from each *Loxosceles* species was reduced with 10 mM DTT and alkylated with 40 mM IAA. Proteins were separated by 10 kDa molecular-weight cut-off ultrafiltration (Microcon YM-10, Millipore, Billerica, MA, USA). The retained proteins (>10kDa) were subsequently digested with sequencing-grade modified trypsin, and the resulting peptides were enriched for glycosylated peptides by hydrophilic interaction liquid chromatography (HILIC), following the procedure described by Alves et al. ^26^.

Glycopeptide intact (non-deglycosylated) samples were analyzed by nano-flow liquid chromatography (EASYnLC, Thermo Fisher Scientific) coupled online to a Q Exactive HF mass spectrometer (Thermo Fisher Scientific). The mass spectrometer was operated in positive ionization mode using data-dependent acquisition (DDA). Full MS scans were acquired in the Orbitrap over an m/z range of 400–1600 at a resolution of 120,000 FWHM, with an automatic gain control (AGC) target of 3 × 10⁶ and a maximum injection time of 100 ms. Following each full MS scan, the 10 most intense precursor ions exceeding an intensity threshold of 1 × 10⁴ were selected for fragmentation. Precursor ions were isolated using a quadrupole isolation window of 1.2 m/z and fragmented by higher-energy collisional dissociation (HCD) using a normalized collision energy of 30. MS/MS spectra were acquired in the Orbitrap at a resolution of 30,000 FWHM, with an AGC target of 1 × 10⁵ and a maximum injection time of 150 ms. Fragmented precursor ions were dynamically excluded from subsequent MS/MS selection for 20 s.

The raw MS spectra from two technical replicates of intact glycopeptides were processed using Byonic software (v4.4.2, Protein Metrics Inc.) against the same Custom *Loxosceles* spp. Protein Database, search parameters and filtering criteria were identical to those previously described (method 1.4). Reverse hits, contaminants, and redundant identifications were excluded from the final dataset.

Peptides were considered non-redundant based on their complete amino acid sequences; sequences differing in length due to distinct cleavage positions were retained separately, although these differences may reflect incomplete or non-specific tryptic digestion rather than distinct peptide forms in the native venom. The relative abundance of each toxin class was calculated by dividing the number of glycosylated peptides assigned to that class by the total number of identified glycosylated peptides, and expressed as a percentage. Glycopeptides assigned to hypothetical proteins, proteins with undefined function, or lacking significant matches to known protein sequences were grouped as “no annotation”. Only confidently identified glycopeptides were considered for the analysis.

Glycosylated phospholipase D sequences identified in gel spots or crude venoms were aligned with the previously described β1Bi sequence ^27^ using Clustal Omega ^28^ and visualized and edited in Jalview v. 2.11.5.2 ^29^. Sequence KU891951.1 was excluded due to the absence of key conserved catalytic and metal-coordination regions.

### 1.7 Enzyme-linked lectin assay (ELLA)

A 96-well polystyrene plates (Corning) were coated overnight at room temperature (RT) with 2 μg/mL of the samples of *L. intermedia*, *L. laeta* and *L. gaucho* venoms, as well as the positive control (fetuin) and negative control (bovine serum albumin -BSA) diluted in PBS pH 7.4. After washing the plates four times with a PBS-Tween 20 0.01%, the plates were blocked with Carbo-Free Blocking Solution (Vector Laboratories) for 30 minutes at RT. Subsequently, the plates were washed four times with wash buffer and the lectins from the biotinylated Lectin Kit I (Vector Laboratories): Concanavalin A (ConA), Soybean agglutinin (SBA), Ricinus communis agglutinin (RCA), Dolichos biflorus agglutinin (DBA), peanut agglutinin (PNA), wheat germ agglutinin (WGA), and Ulex europaeus agglutinin (UEA), were added at a dilution of 1:5000 and the lectins Sambucus nigra agglutinin (SNA), Maackia amurensis lectin II (MAL II) (Vector Laboratories) at 1:10000 dilutions in 1x PBS. The plate was incubated at RT for 1 hour. Afterwards, it was washed 4 times and then incubated with streptavidin-horseradish peroxidase (HRP) (Sigma) diluted 1:10000 in PBS-Tween 0.01% for 1 hour at RT. The last 4 washes were performed and followed by development with the TMB Substrate Kit (Thermo Scientific), mixing in equal parts the TMB solution (3,3’, 5,5-tetramethylbenzidine) with the hydrogen peroxide solution (0.02%). The plate was incubated under protection from light for 15 minutes, and reaction was stopped with 100 μL per well of 2M sulfuric acid. Absorbance was measured in the Multiskan SkyHigh Microplate Spectrophotometer (Thermo Scientific) plate reader at 450 nm. All experiments were done in triplicate.

## Results

### 2DE approach identified a heterogeneous glycoprotein profile with species-specific differences among *Loxosceles* venoms

The total proteomic profile of the venom of the three studied species was obtained using Sypro Ruby (Thermo Scientifics), revealing dispersed proteins distributed in a wide range of isoelectric point (pI) and molecular weight (MW), with a higher concentration of protein spots in the 25 to 37 kDa range (Figure 1A). After staining with the Pro-Q ® Emerald 300 Glycoprotein Gel and Blot Stain kit (Thermo Scientifics), several glycosylated protein spots were detected in the venom of all three species across different pI values, and predominantly within the 25-37 kDa range (Figure 1B, 1C).

**Figure 1.**
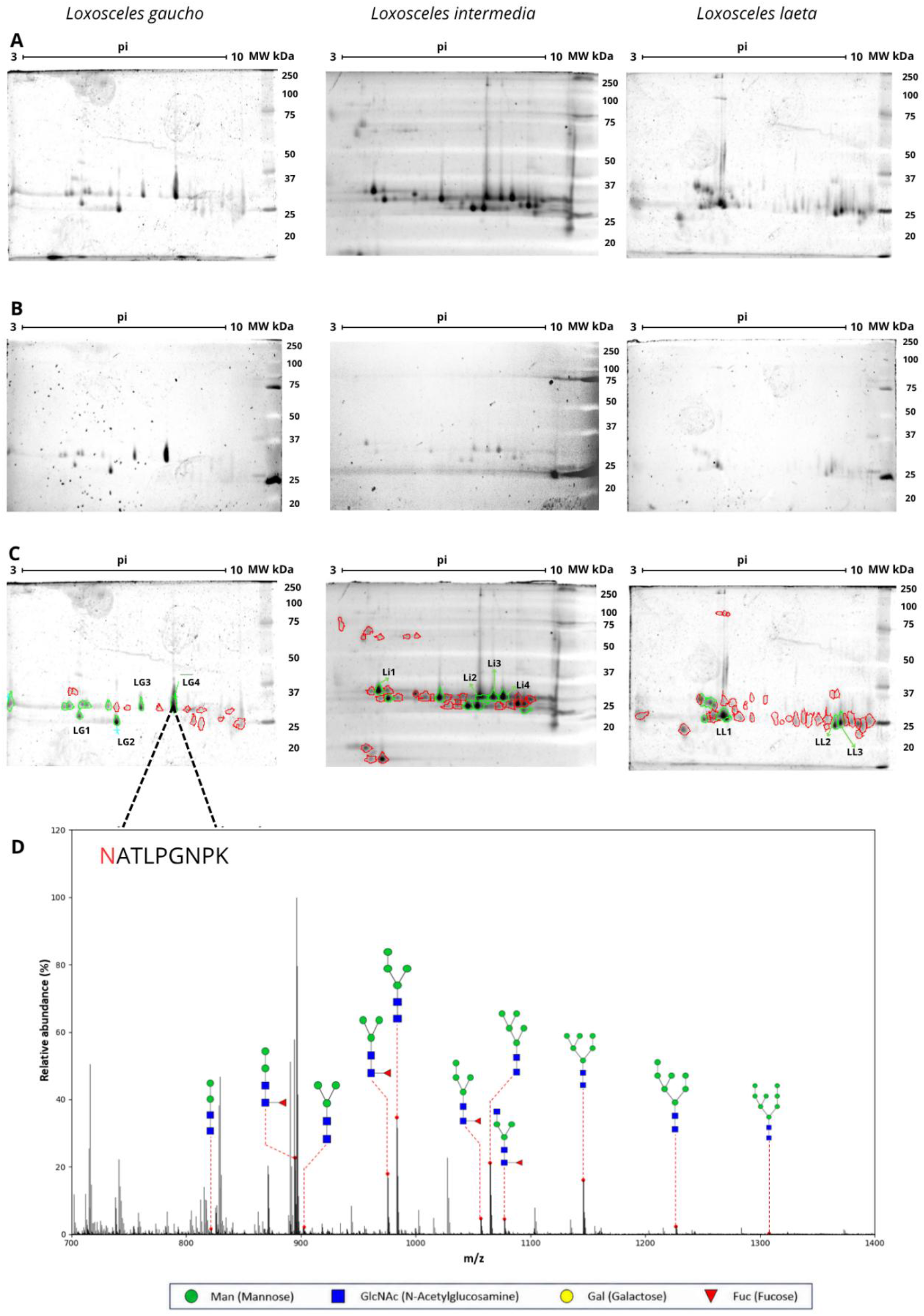
(Glyco)proteomic profile of venoms from spiders of the genus *Loxosceles* obtained by two-dimensional electrophoresis (2DE). Samples containing 60 µg of crude venom from *L. gaucho*, *L. intermedia*, and *L. laeta* were separated by isoelectric focusing (IEF) using 7 cm immobilized pH gradient (IPG) strips with a non-linear pH range of 3–10 to determine the isoelectric point (pI) of the proteins. Subsequently, proteins were resolved by 12.5% SDS-PAGE according to their molecular weight (MW, kDa). (A) SYPRO Ruby staining reveals the total proteomic profile of the venoms from the three species, whereas (B) Pro-Q® Emerald 300 Glycoprotein staining highlights glycosylated proteoforms. (C) Spot detection and image analysis were performed using ImageMaster™ 2D Platinum software. Glycosylated spots are indicated by green circles, and non-glycosylated spots by red circles. (D) Representative MS1 spectrum obtained from the excised gel spot LG4, highlighting the glycan compositions identified by Byonic according to their respective m/z values on the NATLPGNPK peptide sequence. The N-glycosylation site is highlighted in red. The inferred glycan structures corresponding to each assigned composition are schematically illustrated.

Quantitative analysis of the stained gels revealed differences in the proportion of glycosylated proteins among the three species. In the *L. gaucho* venom, 10 glycosylated spots were identified, corresponding to 43.47% of the 23 total protein spots detected with Sypro Ruby staining. *L. intermedia* venom showed 9 glycosylated spots among 36 total protein spots corresponding to 25% of total proteome or 33.33% when only the 27 spots within the 25-37 kDa range were considered. In contrast, *L. laeta* presented the lowest proportion of glycosylated spots identified, with 7 glycosylated spots among 43 total protein spots (16.27%) or 17.5% when considering the 40 spots detected within the 25–37 kDa range (Figure 1C).

To further characterize these glycosylated spots, 11 spots were selected for mass spectrometry analysis, including 4 from *L. intermedia*, 4 from *L. gaucho*, and 3 from *L. laeta*. Glycans were identified after analysis on Byonic software in most of the analyzed spots, except for 2 of the 3 spots from *L. laeta*. As shown in Table S1, all analyzed spots matched different phospholipase D isoforms with high sequence coverage (>60%). In addition, some peptide sequences were recurrently identified as glycosylated across 2 or more spots, particularly “NATLPGNPK” in *L. gaucho* venom and “NATTPGNPK” in *L. intermedia* venom. These sequences were associated with the proteins Loxtox S3A (KU891946.1) and Loxtox I5 (EF535254.1), respectively, both previously described in *Loxosceles* venoms.

The LC–MS/MS analysis also revealed considerable microheterogeneity at individual N-glycosylation sites. In addition to the recurrence of these glycopeptides across different spots, a high diversity of glycan compositions associated with the same N-glycosylation site was observed. One spot exhibited 11 distinct glycan structures associated with a single glycosylation site (Figure 1D). These findings demonstrate the presence of multiple glycan compositions associated with specific phospholipase D proteoforms.

### Identification of diverse N-glycosylated peptides across *Loxosceles* protein families

To expand the glycoproteomic analysis beyond gel-resolved proteins, an additional LC–MS/MS analysis was performed directly on trypsinized crude venom samples. A Custom *Loxosceles* spp. Protein Database was also generated to support peptide identification and glycoproteomic characterization comprising 23,683 *Loxosceles* venom protein sequences.

Using this database, a total of 88 non-redundant N-glycosylated peptide sequences were identified, including 24 from *L. gaucho*, 22 from *L. laeta*, and 42 from *L. intermedia* after processing using Byonic using PEP2D < 0.001 and Score > 100 as a filtering criterion. Supplementary Table 2 summarizes the N-glycosylation analysis of the three venoms, including non-redundant glycosylated peptide sequences, the glycan compositions identified, their glycosylation sites, and the corresponding protein families and classes.

These glycopeptides were distributed across sixteen protein families including several toxins families with established biological activity in *Loxosceles* envenomation, such as phospholipase D, Astacin M12A (Metalloproteases), phospholipase A2, hyaluronidase, S1 protease (serine proteases), among others proteins (Figure 2A). Considering the annotated proteins, glycosylated peptides were predominantly associated with the lyase and hydrolase classes (Figure 2B).

**Figure 2.**
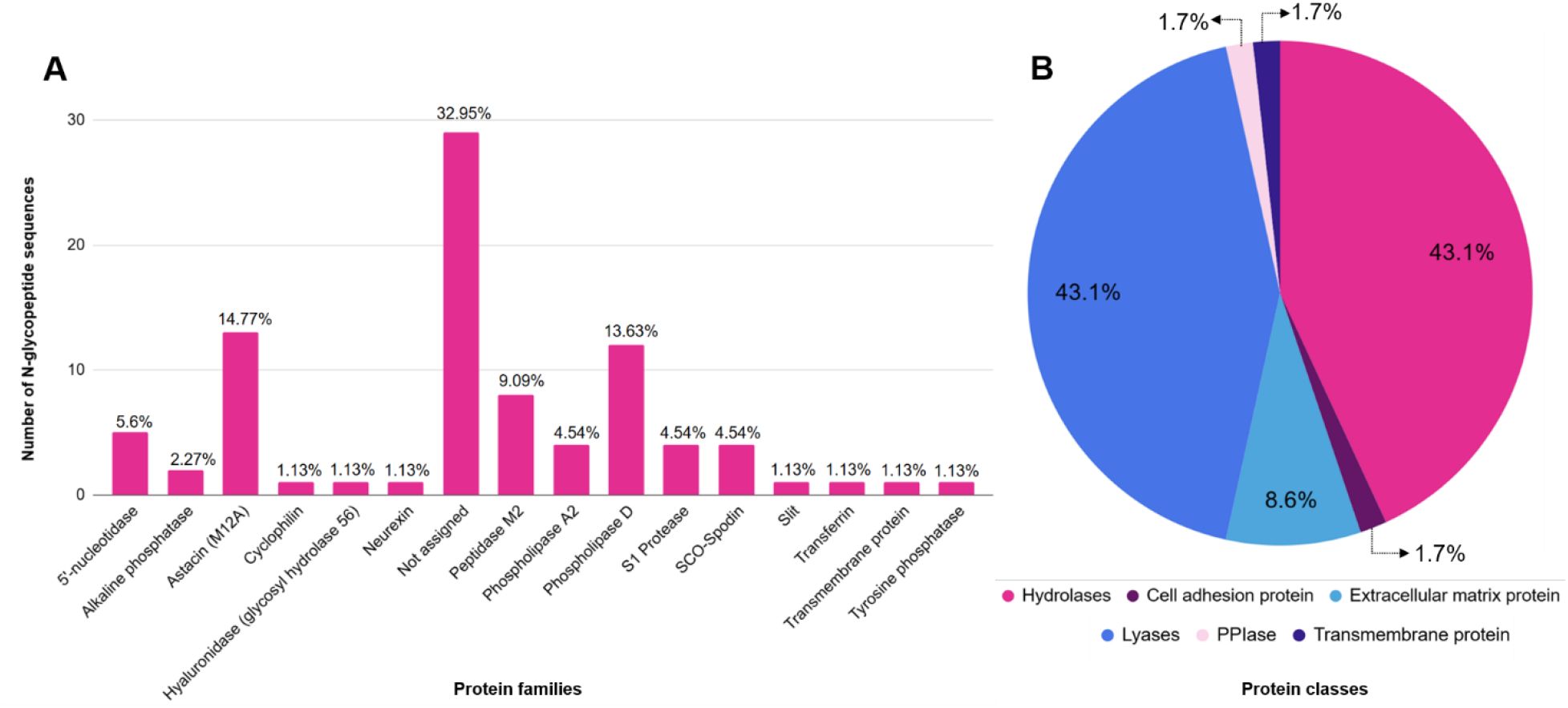
Qualitative analysis of the glycoproteome of *Loxosceles* venom. (A) Distribution of the 88 non-redundant N-glycosylated peptide sequences according to the identified protein families across the three *Loxosceles* species. Bars represent the number of glycosylated peptide sequences assigned to each protein family, while percentages indicate their relative abundance among all identified N-glycopeptides Hypothetical, or uncharacterized proteins and proteins without functional annotation were grouped as “not assigned”. (B) Protein class distribution of the identified glycopeptides.

A substantial proportion of glycosylated peptides (32.95%) corresponded to proteins without functional annotation, hypothetical or uncharacterized proteins. These proteins were classified as proteins for which no protein family or class could be confidently assigned (“not assigned”).

Despite this, glycosylated peptides were widely distributed among biologically relevant protein families and differed markedly among species (Figure S1). The family of the Phospholipases D represented 13.63% of the total glycoproteome, with a higher proportion in *L. gaucho* and *L. laeta* compared to *L. intermedia*. Astacin-like metalloproteases accounted for the largest fraction of glycosylated toxins (14.77%), and were predominantly detected in *L. intermedia*. Glycosylated peptides from 5′-nucleotidase, Angiotesin-converting enzyme, and SCO-spondin were identified in all three species, although with distinct relative abundances, being particularly enriched in *L. laeta*. In contrast, serine proteases and hyaluronidase glycosylated peptides showed species-specific glycosylation patterns, with glycosylated hyaluronidase detected exclusively in *L. intermedia*.

### N-glycopeptide characterization revealed extensive microheterogeneity and a predominance of glycosylation on phospholipase D proteoforms

The glycosylated peptide dataset generated by the crude venom analysis also enabled a detailed characterization of the N-glycan compositions associated with individual glycosylation sites.

As shown in Figure 3, most N-glycosylated peptides presented a single non-redundant glycan composition, and this pattern was observed in the venom of all three analyzed species. Nevertheless, marked microheterogeneity was detected at specific glycosylation sites. For example, in *L. intermedia*, a single sequence exhibited 8, 9, 12, and even 15 different N-glycosylation compositions.

**Figure 3.**
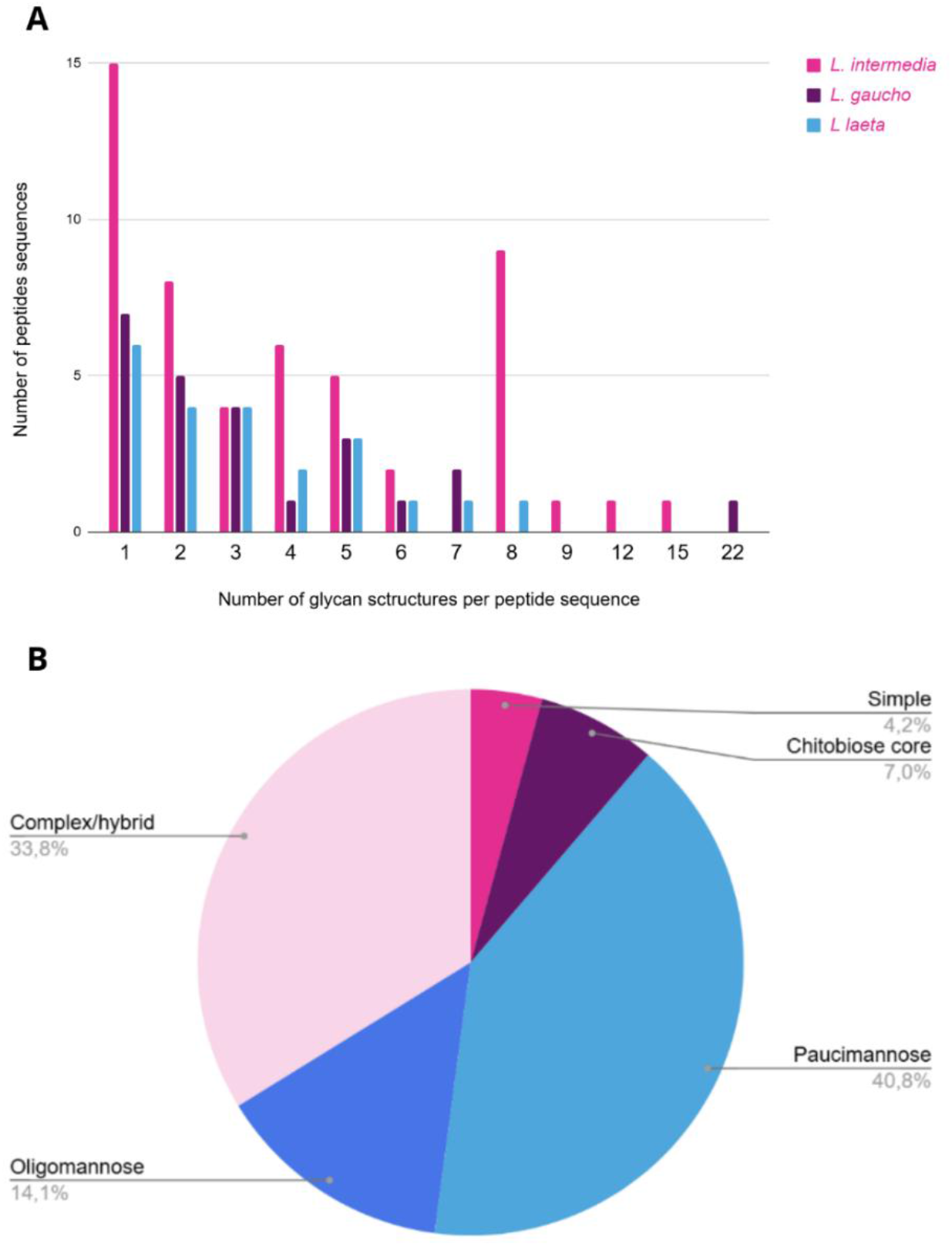
Distribution of N-glycan compositions identified per peptide sequence in *Loxosceles* venoms. (A) Number of glycan compositions detected for each peptide in each species. (B) Distribution of five glycan structure groups (simple, chitobiose core, paucimannose, oligomannose, and complex/hybrid) across the three *Loxosceles* species. Glycan groups were assigned exclusively based on monosaccharide composition. Simple structures were defined as containing a single N-acetylhexosamine (HexNAc), with or without one fucose residue [HexNAc(1) ± Fuc(1)]; chitobiose core structures as HexNAc(2); paucimannose structures as HexNAc(2)Hex(1–3), with or without fucose; oligomannose (high-mannose) structures as HexNAc(2)Hex(4–9); and complex/hybrid structures as compositions containing three or more HexNAc residues [HexNAc(≥3)], with or without additional hexose, fucose, and/or sialic acid residues. Complex and hybrid structures were grouped together because they cannot be reliably distinguished based solely on monosaccharide composition. HexNAc: N-acetylhexosamine; Hex: Hexose; Fuc: Fucose.

The highest degree of glycan microheterogeneity was observed for the peptide NATLPGNPK identified in *L. gaucho* venom, corresponding to the phospholipase D proteoform Loxtox S3A, which was associated with 22 distinct glycoforms (Figure 3A).

To further examine the structural diversity of these glycans, the identified compositions were classified according to their N-glycosylation class. Additionally, structures containing only one HexNAc residue, with or without fucose, were grouped as "simple" structures.

Considering the three species, paucimannose-type glycans represented the largest proportion of identified structures (40.8%), followed by complex/hybrid glycans (33.8%) (Figure 3B). Oligomannose structures were also detected in all three species with similar frequencies. In contrast, simple structures (those containing only one HexNAc residue, with or without fucose) and chitobiose core structures were not detected in *L. laeta* venom and occurred at lower frequencies in *L. gaucho* and *L. intermedia* venom.

Based on the glycan compositions assigned to the identified glycopeptides, GlycoWorkbench was used to infer the corresponding glycan structures (Figure 4). A total of 26 distinct N-glycosylation structures were identified in peptides corresponding to the major toxin families associated with the pathological effects of loxoscelism, including phospholipases D, Astacin-metalloproteases, phospholipase A2, hyaluronidases, and serine proteases (Figure 4). It is important to note that the glycan structures generated using GlycoWorkbench are graphical representations of the identified glycans; carbohydrate linkage information could not be determined in this study.

**Figure 4.**
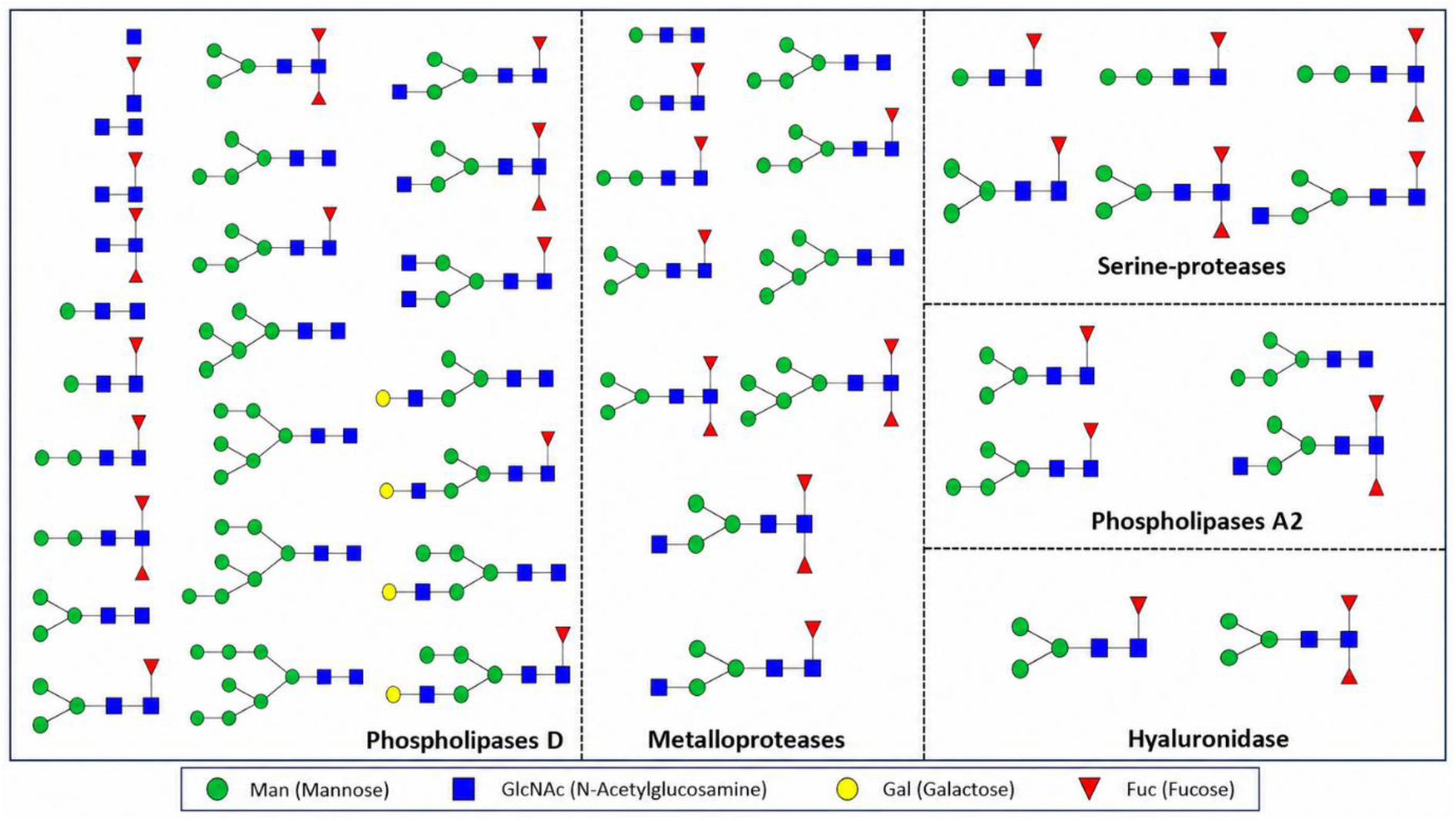
Diversity of glycans identified in peptides belonging to different toxin classes present in the crude venoms of the three analyzed species. Glycan structures were inferred based on glycan compositions obtained through glycoproteomic analysis and represented according to the Symbol Nomenclature for Glycans (SNFG). A total of 26 distinct glycan compositions were identified, predominantly associated with peptides corresponding to phospholipases D. However, glycoforms were also detected in other relevant toxin families, including S1 Proteases (serine proteases), Astacins (metalloproteases), phospholipases A2, and hyaluronidases. The same glycan composition may be associated with different toxin families and/or species, reflecting the shared occurrence of several glycoforms across the analyzed venoms.

All glycan compositions, except HexNAc(2)Hex(5)Fuc(2), were observed in glycopeptides corresponding to phospholipase D proteoforms in at least one species, with most structures being shared among all three species. Additionally, simple structures and chitobiose core structures were detected exclusively in this toxin family. Astacin-like metalloproteases also exhibited substantial structural diversity, with 12 of the 26 identified glycan structures associated with this protein family.

The diversity of glycan compositions identified by LC–MS/MS provided a detailed view of the N-glycosylation patterns in *Loxosceles* venoms. However, mass spectrometry-based approaches may not resolve the full repertoire of glycan structures, particularly when certain glycan features are poorly detected or structurally unresolved.

### Lectin-based characterization of specific carbohydrate epitopes

To further characterize the carbohydrate repertoire of the venoms, lectin-binding assays were therefore employed as a complementary approach, allowing the detection of specific carbohydrate epitopes that may not be readily captured by glycoproteomic analysis alone. Lectin binding was evaluated by enzyme-linked lectin assay (ELLA) using nine lectins types with specificity for mannose (ConA), N-acetylgalactosamine (SBA, DBA), galactose (RCA, PNA), N-acetylglucosamine (WGA), α-fucose (UEA), sialic acid 2-3 (MALL II), and sialic acid 2-6 (SNA).

Lectin-binding profiles (Figure 5) showed consistent reactivity of venoms from all three *Loxosceles* species with ConA and WGA, indicating the presence of mannose and N acetylglucosamine containing glycoconjugates. Species-specific differences were also observed in the binding of lectins recognizing N-acetylgalactosamine and galactose residues (SBA, DBA, RCA, and PNA).

**Figure 5.**
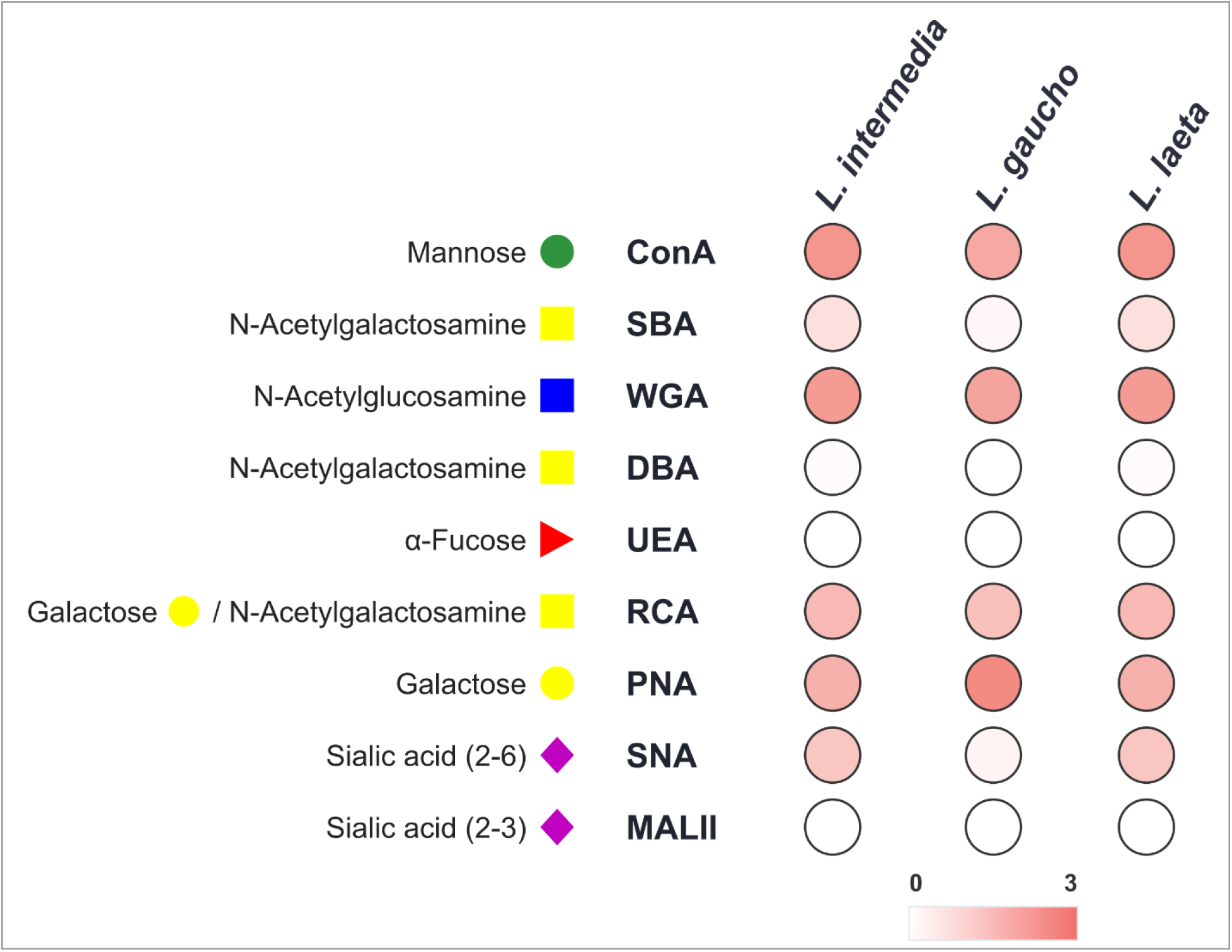
Enzyme-linked lectin assay (ELLA) for carbohydrate detection in *Loxosceles* venoms. Crude venoms (2 µg/mL) were analyzed using biotinylated lectins with distinct carbohydrate specificities: Concanavalin A (ConA), Soybean agglutinin (SBA), Ricinus communis agglutinin (RCA), Dolichos biflorus agglutinin (DBA), Peanut agglutinin (PNA), Wheat germ agglutinin (WGA), Ulex europaeus agglutinin (UEA), Sambucus nigra agglutinin (SNA), and Maackia amurensis lectin II (MAL II). Reactivity is expressed as mean absorbance at 450 nm from triplicate measurements after blank subtraction. Results are presented as a schematic heatmap, with color intensity representing the absorbance values measured by spectrophotometry (0-3). Darker shades indicate higher absorbance values and, consequently, greater lectin reactivity. The same color scale was applied to all lectins. Symbols indicate the carbohydrate specificities of the lectins employed.

Additionally, *L. intermedia* and *L. laeta* venoms showed reactivity with the SNA lectin, specific for α2,6-linked sialic acid, whereas no reactivity was detected with MAL II, specific for specificity for α2,3-linked sialic acid. No reactivity with the α-fucose-specific lectin UEA was detected in any of the venoms, indicating the absence or very low abundance of accessible α-fucose epitopes under the experimental conditions.

## Discussion

The present study provides the first experimental characterization of N-glycosylation in *Loxosceles* venoms through the integration of gel-based glycoproteomic analysis, LC-MS/MS of crude venoms, and lectin profiling. This integrated approach revealed a widespread and structurally heterogeneous N-glycome distributed across multiple venom protein families, including toxins with established roles in envenomation such as phospholipases D, metalloproteases, and hyaluronidases ^30–32^. Several glycosylated peptides could not be confidently assigned to previously characterized proteins, further highlighting the incomplete characterization of the *Loxosceles* venom proteome. These findings indicate that post-translational modifications contribute an additional dimension to venom molecular diversity that is not captured by sequence-based proteomic analyses alone.

N-glycosylation has been reported in toxins from several arthropod groups, including wasps, bees, and scorpions, although the extent of glycan characterization varies among studies. In scorpions and wasps, N-glycosylation has been identified in important venom proteins, although comprehensive structural characterization of their glycan repertoires has not been provided ^25,33^. However, paucimannosidic and hybrid oligosaccharides have been reported in phospholipase A2 (PLA2) from bee venom and phospholipase A1 (PLA1) from ant venom, respectively ^34–36^. In this context, the predominance of paucimannosidic glycans and the frequent occurrence of N-acetylglucosamine, mannose, and core fucose in *Loxosceles* venoms are consistent with glycosylation patterns reported in insects ^37,38^. However, the relatively high occurrence of hybrid and complex structures indicates that the *Loxosceles* venom N-glycome extends beyond the glycan classes commonly associated with insect glycoproteins. Similarly broad structural diversity has been documented in snake venoms: an analysis of eight Bothrops species identified 52 distinct N-glycan compositions corresponding to 60 structures ^39^. Thus, the diversity observed here is consistent with the broad structural variation reported in animal venoms.

The complementary LC-MS/MS and lectin analyses supported the structural diversity of *Loxosceles* venom glycoproteins while highlighting methodological limitations. Lectin assays were broadly consistent with LC-MS/MS for mannose and N-acetylglucosamine, whereas discrepancies were observed for fucose and sialic acid. Fucosylated structures detected by LC-MS/MS were not recognized by UEA-I, possibly because this lectin preferentially binds α-(1,2)-linked fucose, whereas invertebrate N-glycans commonly contain α1,3/1,6-linked core fucose ^40,41^. Conversely, α2,6-sialic acid reactivity was detected in two *Loxosceles* species but was not identified by LC-MS/MS, potentially reflecting the limited glycan library used in the analysis. Although sialylated structures represented a minor fraction of the detected glycans, their biological relevance cannot be excluded, as sialylation occurs in insects and terminal sialic acid has been shown to impact substrate interactions in Naja atra metalloproteinases ^42,43^. Thus, broader glycan libraries and complementary glycomic approaches will be required to determine whether sialylated structures are present in *Loxosceles* venom and whether they have functional consequences.

Multiple N-glycan compositions were also associated with the same peptide sequence, indicating site-specific glycan microheterogeneity. This was particularly evident for PLDs, in which peptides from Loxtox S3A and Loxtox I5 displayed multiple glycan compositions. PLDs already exhibit considerable sequence and isoform diversity ^6,8,9,44,45^ and the glycan heterogeneity identified here suggests an additional source of proteoform variation. Glycan microheterogeneity can alter protein surface properties, including charge, hydration, steric accessibility, and conformational dynamics ^46^, although its functional consequences for *Loxosceles* toxins remain unknown.

The distribution of glycosylation sites among PLDs may have functional implications beyond protein stability and folding. PLD activity depends not only on the catalytic site but also on interactions with lipid substrates at membrane interfaces^27^, which may be modulated by glycosylation. In our analysis, six glycosylation sites across five proteins occurred near the catalytic His-47 ^47^ and closer in primary sequence to the β2α2 loop (PCDCFR), which contributes to lipid interactions and substrate positioning (Figure S2). Additional glycosylation sites were identified near the β6α6 loop (ITNCLP), including one in Loxtox S9A located seven residues from this region. Given the roles of both loops in the lipid-binding interface, these patterns suggest that glycosylation may modulate local structural dynamics or PLD–membrane interactions, potentially influencing substrate accessibility or orientation. However, these functional implications remain to be experimentally validated.

Metalloproteases also showed substantial glycan diversity. These enzymes contribute to extracellular matrix degradation and lesion progression during loxoscelism ^48,49^, making their glycosylation of particular interest. Studies in other venom systems have shown that metalloprotease glycosylation can influence enzymatic activity, substrate interactions, and structural stability ^50–53^. More recently, N-glycan removal from Naja atra metalloproteases altered proteolytic activity and heparin-binding affinity and reduced resistance to chemical and thermal denaturation ^43^. These findings provide a rationale for investigating whether glycan heterogeneity similarly contributes to functional differences among *Loxosceles* metalloprotease proteoforms. At present, however, such effects remain hypothetical.

Several limitations should be considered. First, protein identification and species-specific assignment were limited by the completeness of the sequence database. At the time of database construction, only 52 *L. gaucho* sequences were available, compared with 2,012 for *L. intermedia* and 1,489 for *L. laeta*. Consequently, some *L. gaucho* glycopeptides were assigned to conserved proteins previously described in other *Loxosceles* species. This limitation also highlights the need for improved data availability and annotation in spider venom databases, as both unavailable datasets ^54^ and insufficiently annotated public datasets ^55^ can hinder downstream glycopeptide identification.

The use of 7 cm IPG strips and the enrichment and glycan-library strategies may have limited proteoform resolution and structural glycome coverage. Nevertheless, the concordance between spot-based LC-MS/MS and crude venom glycoproteomics provides independent support for the identified glycoproteins and glycosylation sites, while discrepancies with lectin assays suggest that additional glycan structures may remain undetected. Future studies combining optimized enrichment strategies, broader libraries, and complementary glycomic approaches may expand structural coverage and enable a more comprehensive characterization of the *Loxosceles* N-glycome.

Taken together, these findings establish N-glycosylation as an important component of *Loxosceles* venom molecular diversity, particularly among PLDs and metalloproteases. The occurrence of multiple glycan compositions at individual sites indicates substantial proteoform heterogeneity and suggests that glycosylation may contribute to differences in the physicochemical and functional properties of venom proteins. However, the functional consequences of this heterogeneity remain to be demonstrated experimentally. Because the present study focused exclusively on N-glycosylation, other features, including O-glycosylation, site occupancy, glycan isomerism, linkage, branching, and relative abundance, may further contribute to venom proteoform diversity. These findings provide a foundation for future studies investigating how glycan composition shapes the biochemical and biological properties of *Loxosceles* venom proteins.

## Conclusion

The glycoproteomic characterization of *Loxosceles* venoms revealed a broad diversity of N-glycosylations, including multiple glycoforms associated with medically relevant toxins involved in envenomation. By integrating gel-based glycoproteomic, LC-MS/MS analysis of crude venoms, and lectin-binding assays, this study provides the first comprehensive experimental overview of N-glycosylation in *Loxosceles* venoms. These findings expand the current knowledge of the molecular complexity of spider venoms and establish a framework for future studies investigating the structural and functional significance of glycosylation in these venom toxins.

## Support information

**Table S1.**
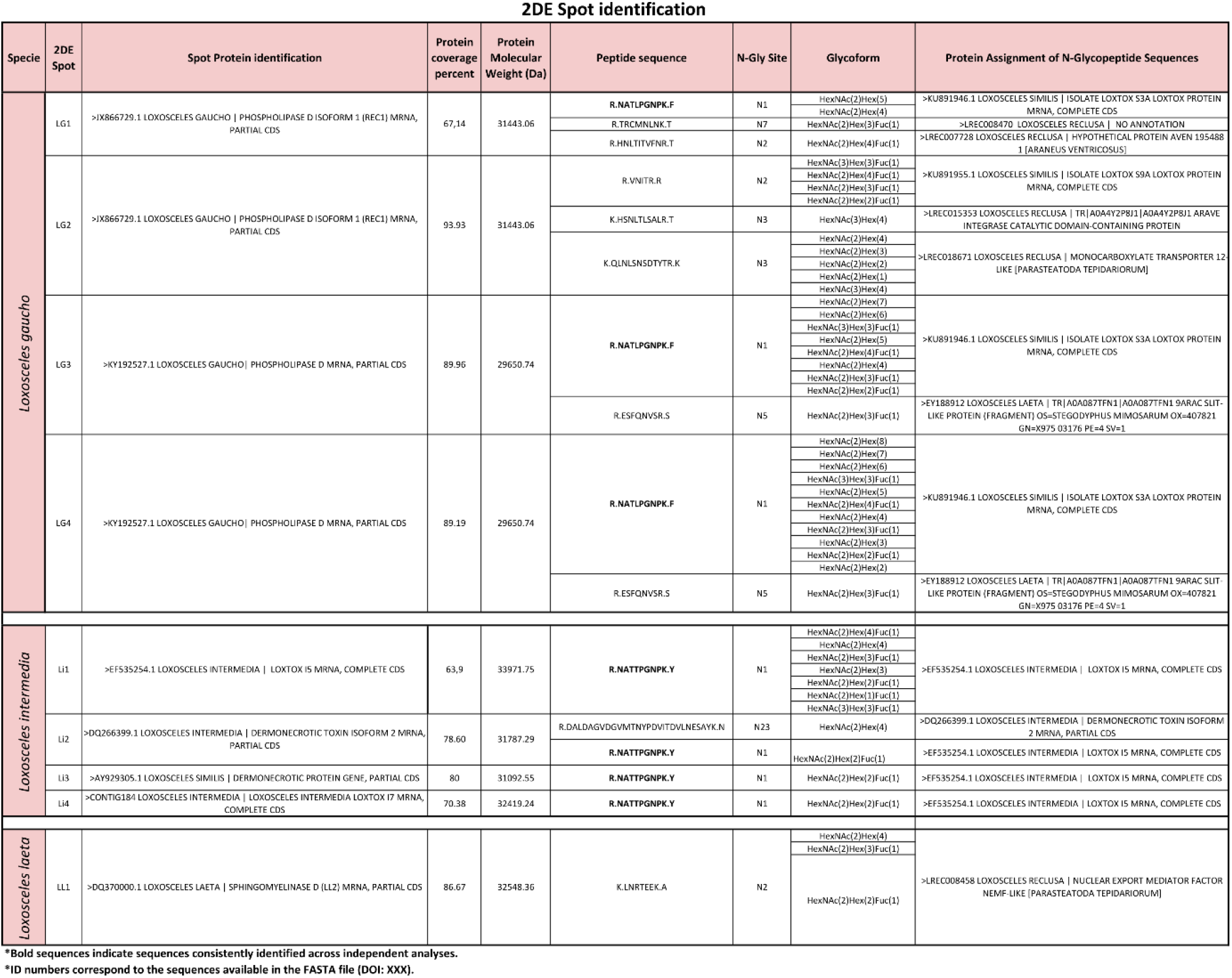
Protein identification and associated N-glycoforms in excised gel spots, including sequence coverage, number of N-glycosylated peptide sequences, glycosylation sites, and glycoform compositions assigned to each site.

**Table S2.**
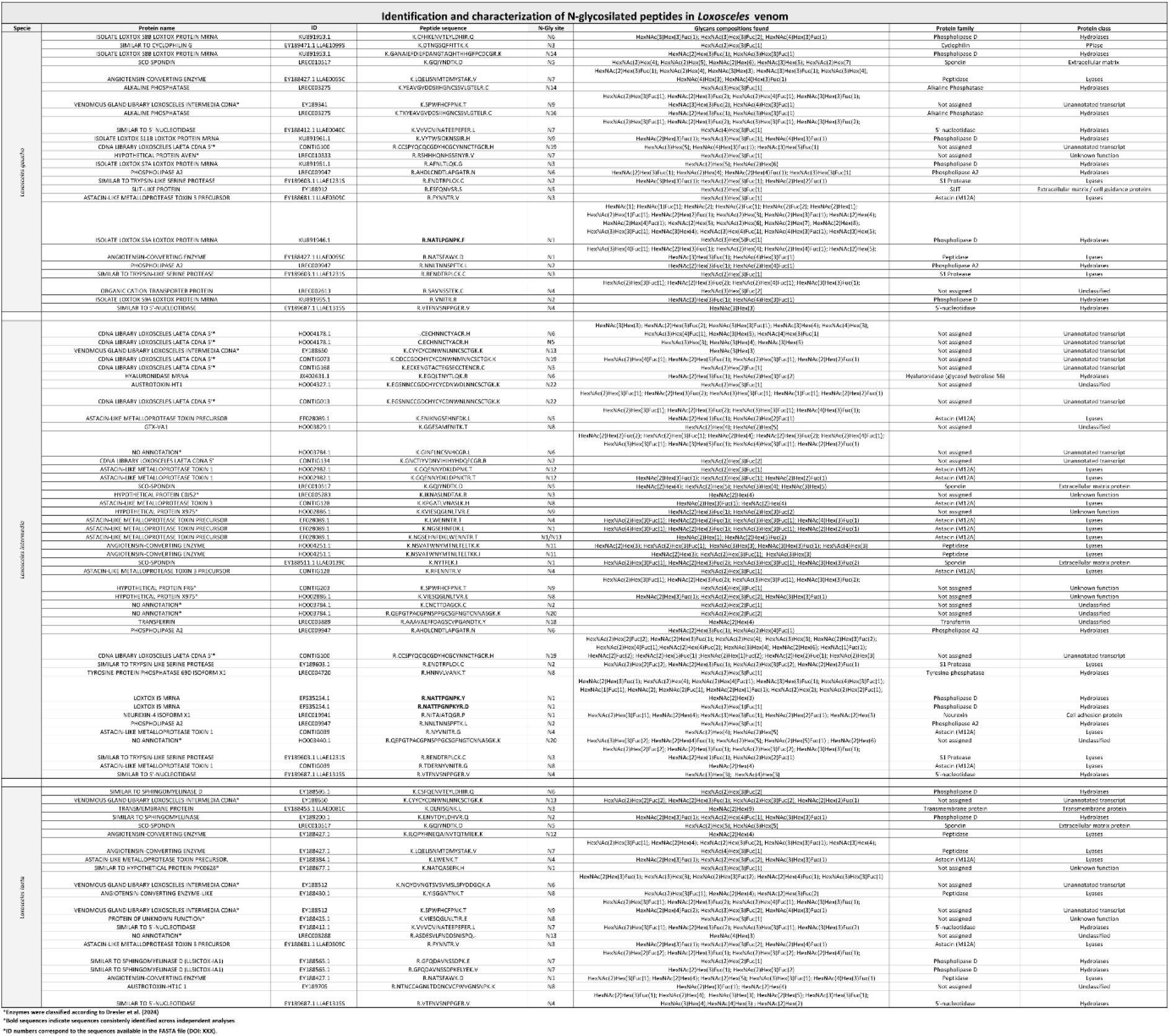
Non-redundant N-glycosylated peptides identified in *Loxosceles* venoms, including peptide sequences, glycosylation sites, and assigned proteins for each species.

**Figure S1.**
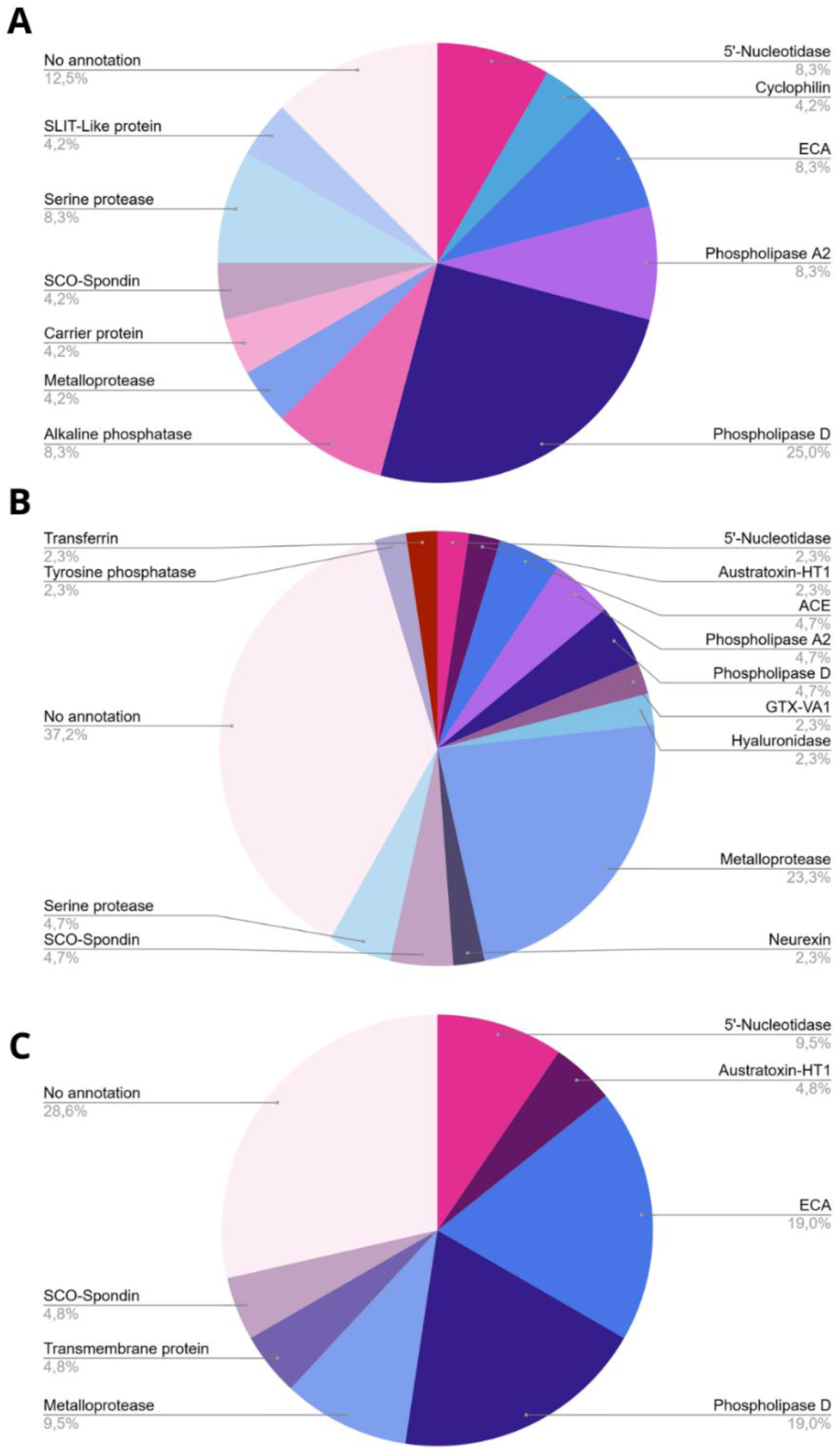
Distribution of N-glycosylated peptides among the protein classes identified in the venoms of (A) *Loxosceles gaucho* (B) *Loxosceles intermedia*, and (C) *Loxosceles laeta*. The relative abundance of glycosylated peptides varied among species, with *L. gaucho* venom showing the highest proportion of N-glycosylated peptides associated with phospholipase D toxins.

**Figure S2.**
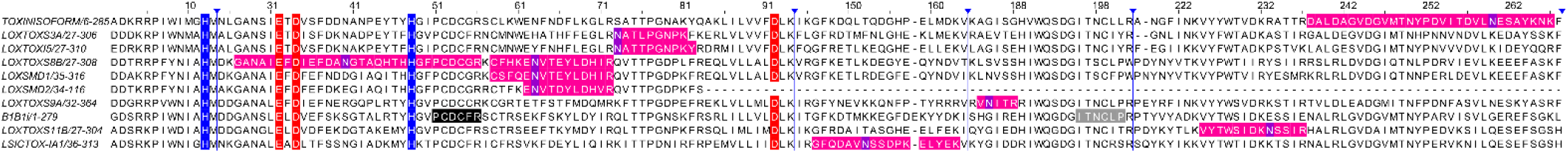
Multiple sequence alignment of N-glycosylated phospholipase D (PLD) sequences and β1B1i. Identified peptides are highlighted in pink, with the N-glycosylation site indicated in purple. Conserved catalytic histidine residues (H12 and H47) are outlined in blue, whereas residues involved in Mg²⁺ coordination are outlined in red. The sequence corresponding to the catalytic β2α2 loop is shown in black, while the flexible loop is shown in gray. The alignment includes the following sequences retrieved from the customized protein database and the NCBI Protein database: LOXTOXS3A (KU891946.1), LOXTOXS9A (KU891955.1), LOXTOXI5 (EF535254.1), TOXINISOFORM2 (DQ266399.1), LOXTOXS8B (KU891953.1), LOXTOXS11B (KU891961.1), LOXTOXS7A (KU891951.1), LOXSMD1 (EY188595.1), LOXSMD2 (EY189200.1), LSICTOX-IA1 (EY188565.1), and β1B1i (A0A0D4WV12).

## Author Contributions

Author contributions: Conceptualization: RSM‡, BCSS‡, MNMB, LF. Methodology: RSM, BCSS, MNMB, MBC, MRL, CCO, HMA, LM, JM. Formal analysis: RSM, DEB, MCEF, MBC Investigation: RSM, BCSS, LM, FM, MNR. Validation: RSM, MNMB, MCEF, CTNR, LF. Data curation: RSM, BCSS, DEB, MCB, MNMB. Writing – original draft: RSM LF. Writing – review and editing: All authors. Funding acquisition: LF. Supervision: LF‡, MNMB‡. Project administration: LF and CCO.

## ACKNOWLEDGMENT

The authors are grateful to the Center for Research and Production of Immunobiologicals (CPPI) for providing the samples used in this study.

