## Supplementary material for "The Glycoproteome of *Loxosceles* Venom Reveals an Expanded Landscape of Animal Toxins": Table_S1

### 2DE Spot identification

<table>
<thead>
<tr><th>Specie</th><th>2DE Spot</th><th>Spot Protein identification</th><th>Protein coverage percent</th><th>Protein Molecular Weight (Da)</th><th>Peptide sequence</th><th>N-Gly Site</th><th>Glycoform</th><th>Protein Assignment of N-Glycopeptide Sequences</th></tr>
</thead>
<tbody>
<tr><td rowspan="35">Loxosceles gaucho</td><td rowspan="4">LG1</td><td rowspan="4">>JX866729.1 LOXOSCELES GAUCHO | PHOSPHOLIPASE D ISOFORM 1 (REC1) MRNA, PARTIAL CDS</td><td rowspan="4">67,14</td><td rowspan="4">31443.06</td><td rowspan="2">R.NATLPGNPK.F</td><td rowspan="2">N1</td><td>HexNAc(2)Hex(5)</td><td rowspan="2">>KU891946.1 LOXOSCELES SIMILIS | ISOLATE LOXTOX S3A LOXTOX PROTEIN MRNA, COMPLETE CDS</td></tr>
<tr><td>HexNAc(2)Hex(4)</td></tr>
<tr><td>R.TRCMNLNK.T</td><td>N7</td><td>HexNAc(2)Hex(3)Fuc(1)</td><td>>LREC008470 LOXOSCELES RECLUSA | NO ANNOTATION</td></tr>
<tr><td>R.HNLTITVFNR.T</td><td>N2</td><td>HexNAc(2)Hex(4)Fuc(1)</td><td>>LREC007728 LOXOSCELES RECLUSA | HYPOTHETICAL PROTEIN AVEN 195488 1 [ARANEUS VENTRICOSUS]</td></tr>
<tr><td rowspan="10">LG2</td><td rowspan="10">>JX866729.1 LOXOSCELES GAUCHO | PHOSPHOLIPASE D ISOFORM 1 (REC1) MRNA, PARTIAL CDS</td><td rowspan="10">93.93</td><td rowspan="10">31443.06</td><td rowspan="4">R.VNITR.R</td><td rowspan="4">N2</td><td>HexNAc(3)Hex(3)Fuc(1)</td><td rowspan="4">>KU891955.1 LOXOSCELES SIMILIS | ISOLATE LOXTOX S9A LOXTOX PROTEIN MRNA, COMPLETE CDS</td></tr>
<tr><td>HexNAc(2)Hex(4)Fuc(1)</td></tr>
<tr><td>HexNAc(2)Hex(3)Fuc(1)</td></tr>
<tr><td>HexNAc(2)Hex(2)Fuc(1)</td></tr>
<tr><td>K.HSNLTLSALR.T</td><td>N3</td><td>HexNAc(3)Hex(4)</td><td>>LREC015353 LOXOSCELES RECLUSA | TR|A0A4Y2P8J1|A0A4Y2P8J1 ARAVE INTEGRASE CATALYTIC DOMAIN-CONTAINING PROTEIN</td></tr>
<tr><td rowspan="5">K.QLNLSNSDTYTR.K</td><td rowspan="5">N3</td><td>HexNAc(2)Hex(4)</td><td rowspan="5">>LREC018671 LOXOSCELES RECLUSA | MONOCARBOXYLATE TRANSPORTER 12-LIKE [PARASTEATODA TEPIDARIORUM]</td></tr>
<tr><td>HexNAc(2)Hex(3)</td></tr>
<tr><td>HexNAc(2)Hex(2)</td></tr>
<tr><td>HexNAc(2)Hex(1)</td></tr>
<tr><td>HexNAc(3)Hex(4)</td></tr>
<tr><td rowspan="9">LG3</td><td rowspan="9">>KY192527.1 LOXOSCELES GAUCHO| PHOSPHOLIPASE D MRNA, PARTIAL CDS</td><td rowspan="9">89.96</td><td rowspan="9">29650.74</td><td rowspan="8">R.NATLPGNPK.F</td><td rowspan="8">N1</td><td>HexNAc(2)Hex(7)</td><td rowspan="8">>KU891946.1 LOXOSCELES SIMILIS | ISOLATE LOXTOX S3A LOXTOX PROTEIN MRNA, COMPLETE CDS</td></tr>
<tr><td>HexNAc(2)Hex(6)</td></tr>
<tr><td>HexNAc(3)Hex(3)Fuc(1)</td></tr>
<tr><td>HexNAc(2)Hex(5)</td></tr>
<tr><td>HexNAc(2)Hex(4)Fuc(1)</td></tr>
<tr><td>HexNAc(2)Hex(4)</td></tr>
<tr><td>HexNAc(2)Hex(3)Fuc(1)</td></tr>
<tr><td>HexNAc(2)Hex(2)Fuc(1)</td></tr>
<tr><td>R.ESFQNVSR.S</td><td>N5</td><td>HexNAc(2)Hex(3)Fuc(1)</td><td>>EY188912 LOXOSCELES LAETA | TR|A0A087TFN1|A0A087TFN1 9ARAC SLIT-LIKE PROTEIN (FRAGMENT) OS=STEGODYPHUS MIMOSARUM OX=407821 GN=X975 03176 PE=4 SV=1</td></tr>
<tr><td rowspan="12">LG4</td><td rowspan="12">>KY192527.1 LOXOSCELES GAUCHO| PHOSPHOLIPASE D MRNA, PARTIAL CDS</td><td rowspan="12">89.19</td><td rowspan="12">29650.74</td><td rowspan="11">R.NATLPGNPK.F</td><td rowspan="11">N1</td><td>HexNAc(2)Hex(8)</td><td rowspan="11">>KU891946.1 LOXOSCELES SIMILIS | ISOLATE LOXTOX S3A LOXTOX PROTEIN MRNA, COMPLETE CDS</td></tr>
<tr><td>HexNAc(2)Hex(7)</td></tr>
<tr><td>HexNAc(2)Hex(6)</td></tr>
<tr><td>HexNAc(3)Hex(3)Fuc(1)</td></tr>
<tr><td>HexNAc(2)Hex(5)</td></tr>
<tr><td>HexNAc(2)Hex(4)Fuc(1)</td></tr>
<tr><td>HexNAc(2)Hex(4)</td></tr>
<tr><td>HexNAc(2)Hex(3)Fuc(1)</td></tr>
<tr><td>HexNAc(2)Hex(3)</td></tr>
<tr><td>HexNAc(2)Hex(2)Fuc(1)</td></tr>
<tr><td>HexNAc(2)Hex(2)</td></tr>
<tr><td>R.ESFQNVSR.S</td><td>N5</td><td>HexNAc(2)Hex(3)Fuc(1)</td><td>>EY188912 LOXOSCELES LAETA | TR|A0A087TFN1|A0A087TFN1 9ARAC SLIT-LIKE PROTEIN (FRAGMENT) OS=STEGODYPHUS MIMOSARUM OX=407821 GN=X975 03176 PE=4 SV=1</td></tr>
<tr><td colspan="9"></td></tr>
<tr><td rowspan="11">Loxosceles intermedia</td><td rowspan="7">Li1</td><td rowspan="7">>EF535254.1 LOXOSCELES INTERMEDIA | LOXTOX I5 MRNA, COMPLETE CDS</td><td rowspan="7">63,9</td><td rowspan="7">33971.75</td><td rowspan="7">R.NATTPGNPK.Y</td><td rowspan="7">N1</td><td>HexNAc(2)Hex(4)Fuc(1)</td><td rowspan="7">>EF535254.1 LOXOSCELES INTERMEDIA | LOXTOX I5 MRNA, COMPLETE CDS</td></tr>
<tr><td>HexNAc(2)Hex(4)</td></tr>
<tr><td>HexNAc(2)Hex(3)Fuc(1)</td></tr>
<tr><td>HexNAc(2)Hex(3)</td></tr>
<tr><td>HexNAc(2)Hex(2)Fuc(1)</td></tr>
<tr><td>HexNAc(2)Hex(1)Fuc(1)</td></tr>
<tr><td>HexNAc(3)Hex(3)Fuc(1)</td></tr>
<tr><td rowspan="2">Li2</td><td rowspan="2">>DQ266399.1 LOXOSCELES INTERMEDIA | DERMONECROTIC TOXIN ISOFORM 2 MRNA, PARTIAL CDS</td><td rowspan="2">78.60</td><td rowspan="2">31787.29</td><td>R.DALDAGVDGVMTNYPDVITDVLNESAYK.N</td><td>N23</td><td>HexNAc(2)Hex(4)</td><td>>DQ266399.1 LOXOSCELES INTERMEDIA | DERMONECROTIC TOXIN ISOFORM 2 MRNA, PARTIAL CDS</td></tr>
<tr><td>R.NATTPGNPK.Y</td><td>N1</td><td>HexNAc(2)Hex(2)Fuc(1)</td><td>>EF535254.1 LOXOSCELES INTERMEDIA | LOXTOX I5 MRNA, COMPLETE CDS</td></tr>
<tr><td>Li3</td><td>>AY929305.1 LOXOSCELES SIMILIS | DERMONECROTIC PROTEIN GENE, PARTIAL CDS</td><td>80</td><td>31092.55</td><td>R.NATTPGNPK.Y</td><td>N1</td><td>HexNAc(2)Hex(2)Fuc(1)</td><td>>EF535254.1 LOXOSCELES INTERMEDIA | LOXTOX I5 MRNA, COMPLETE CDS</td></tr>
<tr><td>Li4</td><td>>CONTIG184 LOXOSCELES INTERMEDIA | LOXOSCELES INTERMEDIA LOXTOX I7 MRNA, COMPLETE CDS</td><td>70.38</td><td>32419.24</td><td>R.NATTPGNPK.Y</td><td>N1</td><td>HexNAc(2)Hex(2)Fuc(1)</td><td>>EF535254.1 LOXOSCELES INTERMEDIA | LOXTOX I5 MRNA, COMPLETE CDS</td></tr>
<tr><td colspan="9"></td></tr>
<tr><td rowspan="3">Loxosceles laeta</td><td rowspan="3">LL1</td><td rowspan="3">>DQ370000.1 LOXOSCELES LAETA | SPHINGOMYELINASE D (LL2) MRNA, PARTIAL CDS</td><td rowspan="3">86.67</td><td rowspan="3">32548.36</td><td rowspan="3">K.LNRTEEK.A</td><td rowspan="3">N2</td><td>HexNAc(2)Hex(4)</td><td rowspan="3">>LREC008458 LOXOSCELES RECLUSA | NUCLEAR EXPORT MEDIATOR FACTOR NEMF-LIKE [PARASTEATODA TEPIDARIORUM]</td></tr>
<tr><td>HexNAc(2)Hex(3)Fuc(1)</td></tr>
<tr><td>HexNAc(2)Hex(2)Fuc(1)</td></tr>
</tbody>
</table>

***Bold sequences indicate sequences consistently identified across independent analyses.**
