## Supplementary material for "The Glycoproteome of *Loxosceles* Venom Reveals an Expanded Landscape of Animal Toxins": Table_S2

### Identification and characterization of N-glycosilated peptides in *Loxosceles* venom

| Specie | Protein name | ID | Peptide sequence | N-Gly site | Glycans compositions found | Protein family | Protein class |
|---|---|---|---|---|---|---|---|
| *Loxosceles gaucho* | ISOLATE LOXTOX S8B LOXTOX PROTEIN MRNA | KU891953.1 | K.CFHKENVTEYLDHIR.Q | N6 | HexNAc(3)Hex(3)Fuc(1); HexNAc(3)Hex(3)Fuc(2); HexNAc(4)Hex(3)Fuc(1) | Phospholipase D | Hydrolases |
| | SIMILAR TO CYCLOPHILIN G | EY189471.1 LLAE1099S | K.DTNGSQFFITTK.K | N3 | HexNAc(2)Hex(3)Fuc(1) | | |
| | ISOLATE LOXTOX S8B LOXTOX PROTEIN MRNA | KU891953.1 | K.GANAIEFDIEFDANGTAQHTHHGFPCDCGR.K | N14 | HexNAc(2)Hex(3)Fuc(1); HexNAc(3)Hex(3)Fuc(1) | Phospholipase D | Hydrolases |
| | SCO-SPONDIN | LREC010517 | K.GQIYNDTK.D | N5 | HexNAc(2)Hex(4); HexNAc(2)Hex(5); HexNAc(2)Hex(6); HexNAc(3)Hex(5); HexNAc(2)Hex(7) | Spondin | Extracellular matrix |
| | ANGIOTENSIN-CONVERTING ENZYME | EY188427.1 LLAE0055C | K.LQELISNMTDMYSTAK.V | N7 | HexNAc(2)Hex(3)Fuc(1); HexNAc(2)Hex(4); HexNAc(3)Hex(3); HexNAc(3)Hex(3)Fuc(1); HexNAc(3)Hex(4); HexNAc(4)Hex(3); HexNAc(4)Hex(3)Fuc(1) | Peptidase | Lyases |
| | ALKALINE PHOSPHATASE | LREC003275 | K.YEAVGVDDSIIHGNCSSVLGTELR.C | N14 | HexNAc(3)Hex(3)Fuc(1) | Alkaline Phosphatase | Hydrolases |
| | VENOMOUS GLAND LIBRARY LOXOSCELES INTERMEDIA CDNA* | EY189341 | K.SPWFHCFPNK.T | N9 | HexNAc(2)Hex(3)Fuc(1); HexNAc(2)Hex(3)Fuc(2); HexNAc(2)Hex(4)Fuc(1); HexNAc(3)Hex(3)Fuc(1); HexNAc(3)Hex(3)Fuc(2); HexNAc(4)Hex(3)Fuc(1) | Not assigned | Unannotated transcript |
| | ALKALINE PHOSPHATASE | LREC003275 | K.TKYEAVGVDDSIIHGNCSSVLGTELR.C | N16 | HexNAc(2)Hex(3)Fuc(1); HexNAc(3)Hex(3)Fuc(1) | Alkaline Phosphatase | Hydrolases |
| | SIMILAR TO 5'-NUCLEOTIDASE | EY188412.1 LLAE0040C | K.VVVCNINATEEPEFER.L | N7 | HexNAc(2)Hex(3)Fuc(1); HexNAc(2)Hex(3)Fuc(2); HexNAc(3)Hex(3)Fuc(1); HexNAc(3)Hex(3)Fuc(2); HexNAc(4)Hex(3)Fuc(1) | 5'-nucleotidase | Hydrolases |
| | ISOLATE LOXTOX S11B LOXTOX PROTEIN MRNA | KU891961.1 | K.VYTWSIDKNSSIR.H | N9 | HexNAc(2)Hex(3)Fuc(1); HexNAc(3)Hex(3)Fuc(1); HexNAc(4)Hex(3)Fuc(1) | Phospholipase D | Hydrolases |
| | CDNA LIBRARY LOXOSCELES LAETA CDNA 5'* | CONTIG100 | R.CCSPYQCQCGDYHCGCYNNCTFGCR.H | N19 | HexNAc(3)Hex(5); HexNAc(4)Hex(3)Fuc(1); HexNAc(3)Hex(5)Fuc(1) | Not assigned | Unannotated transcript |
| | HYPOTHETICAL PROTEIN AVEN* | LREC010333 | R.RSHHHQNHSSENYR.V | N7 | HexNAc(2)Hex(3)Fuc(1) | Not assigned | Unknown function |
| | ISOLATE LOXTOX S7A LOXTOX PROTEIN MRNA | KU891951.1 | R.AFNLTLQK.G | N3 | HexNAc(2)Hex(5); HexNAc(2)Hex(6) | Phospholipase D | Hydrolases |
| | PHOSPHOLIPASE A2 | LREC009947 | R.AHDLCNDTLAPGATR.N | N6 | HexNAc(2)Hex(3)Fuc(1); HexNAc(2)Hex(4); HexNAc(2)Hex(4)Fuc(1); HexNAc(3)Hex(3)Fuc(1) | Phospholipase A2 | Hydrolases |
| | SIMILAR TO TRYPSIN-LIKE SERINE PROTEASE | EY189603.1 LLAE1231S | R.ENDTRPLCK.C | N2 | HexNAc(3)Hex(3)Fuc(1); HexNAc(2)Hex(3)Fuc(1); HexNAc(2)Hex(2)Fuc(1) | S1 Protease | Lyases |
| | SLIT-LIKE PROTEIN | EY188912 | R.ESFQNVSR.S | N5 | HexNAc(2)Hex(3)Fuc(1) | SLIT | Extracellular matrix / cell guidance proteins |
| | ASTACIN-LIKE METALLOPROTEASE TOXIN 3 PRECURSOR | EY188681.1 LLAE0309C | R.FYNNTR.V | N3 | HexNAc(3)Hex(3)Fuc(1) | Astacin (M12A) | Lyases |
| | ISOLATE LOXTOX S3A LOXTOX PROTEIN MRNA | KU891946.1 | **R.NATLPGNPK.F** | N1 | HexNAc(1); HexNAc(1)Fuc(1); HexNAc(2); HexNAc(2)Fuc(1); HexNAc(2)Fuc(2); HexNAc(2)Hex(1); HexNAc(2)Hex(1)Fuc(1); HexNAc(2)Hex(2)Fuc(1); HexNAc(2)Hex(3); HexNAc(2)Hex(3)Fuc(1); HexNAc(2)Hex(4); HexNAc(2)Hex(4)Fuc(1); HexNAc(2)Hex(5); HexNAc(2)Hex(6); HexNAc(2)Hex(7); HexNAc(2)Hex(8); HexNAc(3)Hex(3)Fuc(1); HexNAc(3)Hex(4); HexNAc(3)Hex(4)Fuc(1); HexNAc(4)Hex(3)Fuc(1); HexNAc(3)Hex(5); HexNAc(3)Hex(5)Fuc(1) | Phospholipase D | Hydrolases |
| | ANGIOTENSIN-CONVERTING ENZYME | EY188427.1 LLAE0055C | R.NATSFAWK.D | N1 | HexNAc(3)Hex(4)Fuc(1); HexNAc(2)Hex(3)Fuc(1); HexNAc(2)Hex(4); HexNAc(2)Hex(4)Fuc(1); HexNAc(2)Hex(5); HexNAc(3)Hex(3)Fuc(1); HexNAc(4)Hex(3)Fuc(1) | Peptidase | Lyases |
| | PHOSPHOLIPASE A2 | LREC009947 | R.NNLTNNSPFTK.L | N2 | HexNAc(2)Hex(3)Fuc(1); HexNAc(2)Hex(4)Fuc(1) | Phospholipase A2 | Hydrolases |
| | SIMILAR TO TRYPSIN-LIKE SERINE PROTEASE | EY189603.1 LLAE1231S | R.RENDTRPLCK.C | N3 | HexNAc(2)Hex(3)Fuc(1) | S1 Protease | Lyases |
| | ORGANIC CATION TRANSPORTER PROTEIN | LREC002613 | R.SAVNSSTEK.C | N4 | HexNAc(2)Hex(3)Fuc(1); HexNAc(2)Hex(3)Fuc(2); HexNAc(2)Hex(4)Fuc(1); HexNAc(3)Hex(3)Fuc(1); HexNAc(3)Hex(3)Fuc(2) | Not assigned | Unclassified |
| | ISOLATE LOXTOX S9A LOXTOX PROTEIN MRNA | KU891955.1 | R.VNITR.R | N2 | HexNAc(3)Hex(3)Fuc(1); HexNAc(4)Hex(3)Fuc(1) | Phospholipase D | Hydrolases |
| | SIMILAR TO 5'-NUCLEOTIDASE | EY189687.1 LLAE1315S | R.VTFNVSNPPGER.V | N4 | HexNAc(3)Hex(3) | 5'-nucleotidase | Hydrolases |
| *Loxosceles intermedia* | CDNA LIBRARY LOXOSCELES LAETA CDNA 5'* | HO004178.1 | -.CECHNNCTYACR.H | N6 | HexNAc(3)Hex(3); HexNAc(2)Hex(3)Fuc(2); HexNAc(3)Hex(3)Fuc(1); HexNAc(3)Hex(4); HexNAc(4)Hex(3); HexNAc(3)Hex(4)Fuc(1); HexNAc(3)Hex(5); HexNAc(4)Hex(3)Fuc(1) | Not assigned | Unannotated transcript |
| | CDNA LIBRARY LOXOSCELES LAETA CDNA 5'* | HO004178.1 | C.ECHNNCTYACR.H | N5 | HexNAc(3)Hex(3); HexNAc(3)Hex(4); HexNAc(3)Hex(5) | Not assigned | Unannotated transcript |
| | VENOMOUS GLAND LIBRARY LOXOSCELES INTERMEDIA CDNA* | EY188650 | K.CYYCYCDNWNLNNCSCTGK.K | N13 | HexNAc(3)Hex(3) | Not assigned | Unannotated transcript |
| | CDNA LIBRARY LOXOSCELES LAETA CDNA 5'* | CONTIG073 | K.DDCCGDCHYCYCDNWNMNNCSCTGK.K | N19 | HexNAc(2)Hex(4)Fuc(1); HexNAc(2)Hex(5)Fuc(1); HexNAc(2)Hex(3)Fuc(1); HexNAc(2)Hex(2)Fuc(1) | Not assigned | Unannotated transcript |
| | CDNA LIBRARY LOXOSCELES LAETA CDNA 5'* | CONTIG168 | K.ECKENGTACTEGSECCTENCR.C | N5 | HexNAc(2)Hex(2)Fuc(1) | Not assigned | Unannotated transcript |
| | HYALURONIDASE MRNA | JX402631.1 | K.EGQLTNYTLQK.R | N6 | HexNAc(2)Hex(3)Fuc(1); HexNAc(2)Hex(3)Fuc(2) | Hyaluronidase (glycosyl hydrolase 56) | Hydrolases |
| | AUSTROTOXIN-HT1 | HO004327.1 | K.EGSNNCCGDCHYCYCDNWDLNNCSCTGK.K | N22 | HexNAc(2)Hex(3)Fuc(1) | Not assigned | |
| | CDNA LIBRARY LOXOSCELES LAETA CDNA 5'* | CONTIG013 | K.EGSNNCCGDCHYCYCDNWNLNNCSCTGK.K | N22 | HexNAc(2)Hex(3)Fuc(1); HexNAc(2)Hex(3)Fuc(2); HexNAc(3)Hex(3)Fuc(1); HexNAc(1)Fuc(1); HexNAc(2)Hex(2)Fuc(1) | Not assigned | Unannotated transcript |
| | ASTACIN-LIKE METALLOPROTEASE TOXIN PRECURSOR | EF028089.1 | K.ENIKNGSEHNFDK.L | N5 | HexNAc(2)Hex(3)Fuc(1); HexNAc(2)Hex(3)Fuc(2); HexNAc(3)Hex(3)Fuc(1); HexNAc(4)Hex(3)Fuc(1); HexNAc(2)Hex(1)Fuc(1); HexNAc(2)Hex(2)Fuc(1) | Astacin (M12A) | Lyases |
| | GTX-VA1 | HO003829.1 | K.GGESAMFNITK.T | N8 | HexNAc(2)Hex(4); HexNAc(2)Hex(5) | Not assigned | Unclassified |
| | NO ANNOTATION* | HO003784.1 | K.GINFLNCSNHCGR.L | N6 | HexNAc(2)Hex(2)Fuc(2); HexNAc(2)Hex(3)Fuc(1); HexNAc(2)Hex(4); HexNAc(2)Hex(3)Fuc(2); HexNAc(2)Hex(4)Fuc(1); HexNAc(3)Hex(3)Fuc(1); HexNAc(3)Hex(5)Fuc(1); HexNAc(4)Hex(3)Fuc(1); HexNAc(2)Hex(2)Fuc(1) | Not assigned | Unclassified |
| | CDNA LIBRARY LOXOSCELES LAETA CDNA 5' | CONTIG134 | K.GNCTIYVDNVIHIHYHDQFCGR.R | N2 | HexNAc(2)Hex(3)Fuc(2) | Not assigned | Unannotated transcript |
| | ASTACIN-LIKE METALLOPROTEASE TOXIN 1 | HO002982.1 | K.GQENNYDKLDPNK.T | N12 | HexNAc(2)Hex(3)Fuc(1) | Astacin (M12A) | Lyases |
| | ASTACIN-LIKE METALLOPROTEASE TOXIN 1 | HO002982.1 | K.GQENNYDKLDPNKTR.T | N12 | HexNAc(2)Hex(3)Fuc(1); HexNAc(3)Hex(3)Fuc(1); HexNAc(2)Hex(2)Fuc(1) | Astacin (M12A) | Lyases |
| | SCO-SPONDIN | LREC010517 | K.GQIYNDTK.D | N5 | HexNAc(2)Hex(4); HexNAc(2)Hex(5); HexNAc(3)Hex(4); HexNAc(3)Hex(5) | Spondin | Extracellular matrix protein |
| | HYPOTHETICAL PROTEIN C0J52* | LREC005283 | K.IKNASLNDTAK.R | N3 | HexNAc(2)Hex(4) | Not assigned | Unknown function |
| | ASTACIN-LIKE METALLOPROTEASE TOXIN 3 | CONTIG128 | K.KPGATLVNASLK.H | N8 | HexNAc(2)Hex(3)Fuc(1); HexNAc(2)Hex(4) | Astacin (M12A) | Lyases |
| | HYPOTHETICAL PROTEIN X975* | HO002886.1 | K.KVIESQGLNLTVR.E | N9 | HexNAc(2)Hex(3)Fuc(1); HexNAc(2)Hex(3)Fuc(2) | Not assigned | Unknown function |
| | ASTACIN-LIKE METALLOPROTEASE TOXIN PRECURSOR | EF028089.1 | K.LWENNTR.T | N4 | HexNAc(2)Hex(3)Fuc(1); HexNAc(2)Hex(2)Fuc(1); HexNAc(3)Hex(3)Fuc(1); HexNAc(4)Hex(3)Fuc(1) | Astacin (M12A) | Lyases |
| | ASTACIN-LIKE METALLOPROTEASE TOXIN PRECURSOR | EF028089.1 | K.NGSEHNFDK.L | N1 | HexNAc(4)Hex(3)Fuc(1); HexNAc(2)Hex(3)Fuc(1); HexNAc(3)Hex(3)Fuc(1); HexNAc(2)Hex(2)Fuc(1) | Astacin (M12A) | Lyases |
| | ASTACIN-LIKE METALLOPROTEASE TOXIN PRECURSOR | EF028089.1 | K.NGSEHNFDKLWENNTR.T | N1/N13 | HexNAc(2)Hex(1); HexNAc(2)Hex(5)Fuc(2) | Astacin (M12A) | Lyases |
| | ANGIOTENSIN-CONVERTING ENZYME | HO004251.1 | K.NSVATWNYMTNLTEETK.K | N11 | HexNAc(2)Hex(3); HexNAc(2)Hex(3)Fuc(1); HexNAc(3)Hex(3); HexNAc(3)Hex(3)Fuc(1); HexNAc(4)Hex(3) | Peptidase | Lyases |
| | ANGIOTENSIN-CONVERTING ENZYME | HO004251.1 | K.NSVATWNYMTNLTEETKK.I | N11 | HexNAc(2)Hex(3); HexNAc(2)Hex(3)Fuc(1); HexNAc(3)Hex(3) | Peptidase | Lyases |
| | SCO-SPONDIN | EY188511.1 LLAE0139C | K.NYTFEK.I | N1 | HexNAc(2)Hex(3)Fuc(1); HexNAc(2)Hex(3)Fuc(2); HexNAc(3)Hex(3)Fuc(1); HexNAc(3)Hex(3)Fuc(2) | Spondin | Extracellular matrix protein |
| | ASTACIN-LIKE METALLOPROTEASE TOXIN 3 PRECURSOR | CONTIG128 | K.RFENNTR.V | N4 | HexNAc(2)Hex(3)Fuc(1) | Astacin (M12A) | Lyases |
| | HYPOTHETICAL PROTEIN FR6* | CONTIG203 | K.SPWFHCFPNK.T | N9 | HexNAc(2)Hex(3)Fuc(1); HexNAc(2)Hex(3)Fuc(2); HexNAc(3)Hex(3)Fuc(1); HexNAc(3)Hex(3)Fuc(2); HexNAc(4)Hex(3)Fuc(1) | Not assigned | Unknown function |
| | HYPOTHETICAL PROTEIN X975* | HO002886.1 | K.VIESQGLNLTVR.E | N8 | HexNAc(2)Hex(3)Fuc(1); HexNAc(2)Hex(3)Fuc(2); HexNAc(3)Hex(3)Fuc(1) | Not assigned | Unknown function |
| | NO ANNOTATION* | HO003784.1 | K.CNCTTDAGCK.C | N2 | HexNAc(2)Hex(2)Fuc(1) | Not assigned | Unclassified |
| | NO ANNOTATION* | HO003784.1 | R.QEPGTPACGPNSPPGCSGFNGTCNNASGK.K | N20 | HexNAc(2)Hex(2)Fuc(2) | Not assigned | Unclassified |
| | TRANSFERRIN | LREC003889 | R.AAAVAEFFDAGSCVPGANDTK.Y | N18 | HexNAc(2)Hex(4) | Transferrin | |
| | PHOSPHOLIPASE A2 | LREC009947 | R.AHDLCNDTLAPGATR.N | N6 | HexNAc(2)Hex(3)Fuc(1); HexNAc(2)Hex(4)Fuc(1) | Phospholipase A2 | Hydrolases |
| | CDNA LIBRARY LOXOSCELES LAETA CDNA 5'* | CONTIG100 | R.CCSPYQCQCGDYHCGCYNNCTFGCR.H | N19 | HexNAc(2)Hex(2)Fuc(2); HexNAc(2)Hex(3)Fuc(1); HexNAc(2)Hex(4); HexNAc(3)Hex(3); HexNAc(2)Hex(3)Fuc(2); HexNAc(2)Hex(4)Fuc(1);HexNAc(2)Hex(4)Fuc(2); HexNAc(3)Hex(4); HexNAc(2)Hex(6); HexNAc(1)Fuc(1); HexNAc(2)Fuc(2); HexNAc(2)Hex(5)Fuc(1) ;HexNAc(2)Hex(1)Fuc(2); HexNAc(2)Hex(2)Fuc(1); HexNAc(2)Hex(3) | Not assigned | Unannotated transcript |
| | SIMILAR TO TRYPSIN-LIKE SERINE PROTEASE | EY189603.1 | R.ENDTRPLCK.C | N2 | HexNAc(2)Hex(2)Fuc(2); HexNAc(2)Hex(3)Fuc(1); HexNAc(2)Hex(3)Fuc(2); HexNAc(2)Hex(2)Fuc(1) | S1 Protease | Lyases |
| | TYROSINE-PROTEIN PHOSPHATASE 69D ISOFORM X1 | LREC004720 | R.HNNVLVANK.T | N8 | HexNAc(2)Hex(3)Fuc(1) | Tyrosine-phosphatase | Hydrolases |
| | LOXTOX I5 MRNA | EF535254.1 | **R.NATTPGNPK.Y** | N1 | HexNAc(2)Hex(3)Fuc(1); HexNAc(2)Hex(4); HexNAc(2)Hex(4)Fuc(1); HexNAc(3)Hex(3)Fuc(1); HexNAc(4)Hex(3)Fuc(1); HexNAc(1)Fuc(1); HexNAc(2); HexNAc(2)Fuc(1); HexNAc(2)Hex(1)Fuc(1); HexNAc(2)Hex(2); HexNAc(2)Hex(2)Fuc(1); HexNAc(2)Hex(3) | Phospholipase D | Hydrolases |
| | LOXTOX I5 MRNA | EF535254.1 | **R.NATTPGNPKYR.D** | N1 | HexNAc(2)Hex(1)Fuc(1) | Phospholipase D | Hydrolases |
| | NEUREXIN-4 ISOFORM X1 | LREC019941 | R.NITAIATQGR.P | N1 | HexNAc(2)Hex(3)Fuc(1); HexNAc(2)Hex(4); HexNAc(3)Hex(3)Fuc(1); HexNAc(2)Hex(2)Fuc(1); HexNAc(2)Hex(3) | Neurexin | Cell adhesion protein |
| | PHOSPHOLIPASE A2 | LREC009947 | R.NNLTNNSPFTK.L | N2 | HexNAc(2)Hex(3)Fuc(1) | Phospholipase A2 | Hydrolases |
| | ASTACIN-LIKE METALLOPROTEASE TOXIN 1 | CONTIG039 | R.NYVNITR.G | N4 | HexNAc(2)Hex(4); HexNAc(2)Hex(5) | Astacin (M12A) | Lyases |
| | NO ANNOTATION* | HO003440.1 | R.QEPGTPACGPNSPPGCSGFNGTCNNASGK.K | N20 | HexNAc(3)Hex(3)Fuc(2); HexNAc(2)Hex(4)Fuc(1); HexNAc(2)Hex(5); HexNAc(2)Hex(5)Fuc(1) ; HexNAc(2)Hex(6) | Not assigned | Unclassified |
| | SIMILAR TO TRYPSIN-LIKE SERINE PROTEASE | EY189603.1 LLAE1231S | R.RENDTRPLCK.C | N3 | HexNAc(2)Hex(2)Fuc(2); HexNAc(2)Hex(3)Fuc(1); HexNAc(2)Hex(3)Fuc(2); HexNAc(3)Hex(3)Fuc(1); HexNAc(2)Hex(1)Fuc(1); HexNAc(2)Hex(2)Fuc(1) | S1 Protease | Lyases |
| | ASTACIN-LIKE METALLOPROTEASE TOXIN 1 | CONTIG039 | R.TDERNYVNITR.G | N8 | HexNAc(2)Hex(4) | Astacin (M12A) | Lyases |
| | SIMILAR TO 5'-NUCLEOTIDASE | EY189687.1 LLAE1315S | R.VTFNVSNPPGER.V | N4 | HexNAc(3)Hex(3); HexNAc(4)Hex(3) | 5'-nucleotidase | Hydrolases |
| *Loxosceles laeta* | SIMILAR TO SPHINGOMYELINASE D | EY188595.1 | K.CSFQENVTEYLDHIR.Q | N6 | HexNAc(2)Hex(3)Fuc(2) | Phospholipase D | Hydrolases |
| | VENOMOUS GLAND LIBRARY LOXOSCELES INTERMEDIA CDNA* | EY188650 | K.CYYCYCDNWNLNNCSCTGK.K | N13 | HexNAc(2)Hex(2)Fuc(2); HexNAc(2)Hex(3)Fuc(1); HexNAc(2)Hex(3)Fuc(2); HexNAc(2)Hex(2)Fuc(1) | Not assigned | Unannotated transcript |
| | TRANSMEMBRANE PROTEIN | EY188453.1 LLAE0081C | K.DLNISGNK.L | N3 | HexNAc(2)Hex(9) | Transmembrane protein | Transmembrane protein |
| | SIMILAR TO SPHINGOMYELINASE | EY189200.1 | K.ENVTDYLDHVR.Q | N2 | HexNAc(2)Hex(3)Fuc(1); HexNAc(2)Hex(4)Fuc(1); HexNAc(3)Hex(3)Fuc(1) | Phospholipase D | Hydrolases |
| | SCO-SPONDIN | LREC010517 | K.GQIYNDTK.D | N5 | HexNAc(2)Hex(5); HexNAc(3)Hex(5) | Spondin | Extracellular matrix protein |
| | ANGIOTENSIN-CONVERTING ENZYME | EY188427.1 | K.ILQPYHNEQAINVTQTMIEK.K | N12 | HexNAc(2)Hex(4) | Peptidase | Lyases |
| | ANGIOTENSIN-CONVERTING ENZYME | EY188427.1 | K.LQELISNMTDMYSTAK.V | N7 | HexNAc(2)Hex(3)Fuc(1); HexNAc(2)Hex(4); HexNAc(2)Hex(3)Fuc(2); HexNAc(3)Hex(3)Fuc(1); HexNAc(3)Hex(4); HexNAc(4)Hex(3)Fuc(1) | Peptidase | Lyases |
| | ASTACIN-LIKE METALLOPROTEASE TOXIN PRECURSOR. | EY188384.1 | K.LWENK.T | N4 | HexNAc(2)Hex(3)Fuc(1); HexNAc(2)Hex(3)Fuc(2); HexNAc(3)Hex(3)Fuc(1) | Astacin (M12A) | Lyases |
| | SIMILAR TO HYPOTHETICAL PROTEIN PY00628* | EY188677.1 | K.NATQASEFK.H | N1 | HexNAc(3)Hex(3)Fuc(1) | Not assigned | Unknown function |
| | VENOMOUS GLAND LIBRARY LOXOSCELES INTERMEDIA CDNA* | EY188512 | K.NQYDVNGTSVSVMSLSFYDDGQK.A | N6 | HexNAc(2)Hex(3)Fuc(1); HexNAc(3)Hex(3); HexNAc(2)Hex(3)Fuc(2); HexNAc(2)Hex(4)Fuc(1); HexNAc(3)Hex(3)Fuc(1) | Not assigned | Unannotated transcript |
| | ANGIOTENSIN-CONVERTING ENZYME-LIKE | EY188430.1 | K.YISGGNTNK.T | N8 | HexNAc(2)Hex(3)Fuc(1); HexNAc(2)Hex(4); HexNAc(2)Hex(3)Fuc(2) | Peptidase | Lyases |
| | VENOMOUS GLAND LIBRARY LOXOSCELES INTERMEDIA CDNA* | EY188512 | K.SPWFHCFPNK.T | N9 | HexNAc(2)Hex(3)Fuc(1); HexNAc(2)Hex(3)Fuc(2); HexNAc(2)Hex(4)Fuc(1); HexNAc(3)Hex(3)Fuc(1); HexNAc(2)Hex(4)Fuc(2); HexNAc(3)Hex(3)Fuc(2); HexNAc(4)Hex(3)Fuc(1) | Not assigned | Unannotated transcript |
| | PROTEIN OF UNKNOWN FUNCTION* | EY188425.1 | K.VIESQGLNLTIR.E | N8 | HexNAc(2)Hex(3)Fuc(1) | Not assigned | Unknown function |
| | SIMILAR TO 5'-NUCLEOTIDASE | EY188412.1 | K.VVVCNINATEEPEFER.L | N7 | HexNAc(2)Hex(3)Fuc(1); HexNAc(2)Hex(3)Fuc(2); HexNAc(3)Hex(3)Fuc(1); HexNAc(3)Hex(3)Fuc(2) | 5'-nucleotidase | Hydrolases |
| | NO ANNOTATION* | LREC008288 | R.ASDESVLFNDDSNISPQ.- | N13 | HexNAc(4)Hex(3) | Not assigned | Unclassified |
| | ASTACIN-LIKE METALLOPROTEASE TOXIN 3 PRECURSOR | EY188681.1 LLAE0309C | R.FYNNTR.V | N3 | HexNAc(2)Hex(3)Fuc(1); HexNAc(2)Hex(3)Fuc(2); HexNAc(2)Hex(4)Fuc(1) | Astacin (M12A) | Lyases |
| | SIMILAR TO SPHINGOMYELINASE D (LLSICTOX-IA1) | EY188565.1 | R.GFQDAVNSSDPK.E | N7 | HexNAc(2)Hex(2)Fuc(2); HexNAc(2)Hex(3)Fuc(1); HexNAc(2)Hex(3)Fuc(2); HexNAc(2)Hex(4)Fuc(1); HexNAc(2)Hex(2)Fuc(1) | Phospholipase D | Hydrolases |
| | SIMILAR TO SPHINGOMYELINASE D (LLSICTOX-IA1) | EY188565.1 | R.GFQDAVNSSDPKELYEK.V | N7 | HexNAc(2)Hex(3)Fuc(1); HexNAc(2)Hex(3)Fuc(2) | Phospholipase D | Hydrolases |
| | ANGIOTENSIN-CONVERTING ENZYME | EY188427.1 | R.NATSFAWK.D | N1 | HexNAc(2)Hex(3)Fuc(1); HexNAc(2)Hex(4); HexNAc(2)Hex(5); HexNAc(3)Hex(3)Fuc(1); HexNAc(4)Hex(3)Fuc(1) | Peptidase | Lyases |
| | AUSTROTOXIN-HT1C 1 | EY189705 | R.NTNCCAGNLTDDNCVCFWVGNSNPK.K | N8 | HexNAc(2)Hex(3)Fuc(1); HexNAc(2)Hex(4) | Not assigned | |
| | SIMILAR TO 5'-NUCLEOTIDASE | EY189687.1 LLAE1315S | R.VTFNVSNPPGER.V | N4 | HexNAc(2)Hex(3)Fuc(1); HexNAc(2)Hex(4); HexNAc(3)Hex(3); HexNAc(2)Hex(5); HexNAc(3)Hex(3)Fuc(1); HexNAc(3)Hex(4);HexNAc(4)Hex(3) ; HexNAc(2)Hex(2) | 5'-nucleotidase | Hydrolases |

***Enzymes were classified according to Dresler et al. (2024)**

***Bold sequences indicate sequences consistenly identified across independent analyses**
